# Differential medial temporal lobe and default-mode network functional connectivity and morphometric changes in Alzheimer’s Disease

**DOI:** 10.1101/232165

**Authors:** Kamil A. Grajski, Steven L. Bressler, for the Alzheimer’s Disease Neuroimaging Initiative

## Abstract

We report group level differential detection of medial temporal lobe resting-state functional connectivity disruption and morphometric changes in the transition from cognitively normal to early mild cognitive impairment in an age-, education- and gender-matched 105 subjects Alzheimer’s Disease Neuroimaging Initiative dataset. In mild Alzheimer’s Disease, but not early mild cognitive impairment, characteristic brain atrophy was detected in FreeSurfer estimates of cortical thickness and subcortical and hippocampal subfield volumes. By contrast, functional connectivity analysis detected earlier significant changes. In early mild cognitive impairment these changes involved medial temporal lobe regions of transentorhinal, perirhinal and entorhinal cortices (associated with the earliest stages of neurofibrillary changes in Alzheimer’s Disease), hippocampus, parahippocampal gyrus and temporal pole, and cortical regions comprising or co-activated with the default-mode network, including rostral and medial prefrontal cortex, anterior cingulate cortex, precuneus and inferior temporal cortex. Key findings include: a) focal and bilaterally symmetric spatial organization of affected medial temporal lobe regions; b) mutual hyperconnectivity bilaterally involving ventral medial temporal lobe structures (temporal pole, uncus); and c) dorsal medial temporal lobe hypoconnectivity with anterior and posterior midline default-mode network nodes. These findings position medial temporal lobe resting state functional connectivity as a candidate biomarker of an Alzheimer’s Disease pathophysiological cascade, potentially in advance of clinical biomarkers, and coincident with biomarkers of the earliest stages of Alzheimer’s neuropathology. Our results indicate that medial temporal lobe resting-state functional connectivity should be further investigated as a potential biomarker in the diagnosis of Alzheimer’s Disease.

**Highlights:**

- Functional connectivity change seen before structural change in Alzheimer’s Disease
- Medial temporal lobes mutually hyper-connect in mild cognitive impairment
- Medial temporal lobe and default mode network decouple in mild cognitive impairment
- Loci of functional change in hippocampi are focal with bilaterally symmetric features
- Nonmonotonic functional connectivity changes in Alzheimer’s Disease progression

## 1. Introduction

In late-onset Alzheimer’s Disease, neuropathologies may cumulate for decades before possible though not inevitable clinical manifestation. Diagnostic guidelines (Albert, 2011; Jack, 2011a; McKhann, 2011; Sperling, 2011) and a research framework (Jack, et al., 2018) for cognitively normal (CN), mild cognitive impairment (MCI), and Alzheimer’s Disease individuals, consider multiple neuroimaging modalities including amyloid β and tau tracers, markers of metabolic activity (FDG PET), and morphometry (MRI). An hypothesized Alzheimer’s Disease pathophysiological cascade links these neuroimaging modalities together with cerebrospinal fluid (CSF) markers for amyloid β and tau to the Alzheimer’s Disease clinical progression (Jack, 2011b, 2013).

The hypothesis tested in this investigation was that functional connectivity disruption can be detected in the early MCI group. Disruption of functional connectivity in Alzheimer’s Disease has been observed in the default-mode network and other resting-state networks in resting-state functional MRI (rsfMRI) (Greicius, et al., 2004; Buckner, et al., 2005; Sorg, et al., 2007; Sperling, et al., 2010; Zhou, et al., 2010; Jones, et al., 2011; Jones, et al., 2012, 2016; Brier, et al., 2012; Ward, et al., 2014; Cai, et al., 2015; Dillen, et al., 2017). However, such observations have not broadly impacted clinical practice (Woo, et al., 2017).

The purpose of the present study is thus to compare functional biomarkers using functional connectivity analysis of resting-state fMRI, with structural imaging biomarkers serving as a reference baseline. The experimental design is a group-level analysis of an Alzheimer’s Disease Neuroimaging Initiative (ADNI) population containing clinically-defined cognitively normal, early MCI and mild Alzheimer’s dementia subgroups. We observed cortical thinning, subcortical volume loss, and whole hippocampal and hippocampal subfield volume loss, in the mild Alzheimer’s Disease subgroup, but not in the early MCI group. A group-level pairwise region of interest (ROI) analysis was used to assess functional connectivity changes. *A priori*, the first element in each ROI pair (the “anchor ROI”) was in the medial temporal lobe (MTL) identified by a method described below. The reason to anchor each ROI pair in the MTL was that it is in the MTL perirhinal cortex (Brodmann Areas 35 and 36) and, specifically, the transentorhinal region (Brodmann Area 35) that the neurofibrillary changes associated with Alzheimer’s Disease start and then, in a progression with disease stages, appear in entorhinal cortex (Brodmann Area 28), hippocampus and temporal cortex, association cortices, and ultimately primary sensory cortices (Braak and Braak, 1985, 1991). The ROI position of the second element in each identified ROI pair was determined in a data-driven method described below. The combination of *a priori* and data-driven approach directly linked functional connectivity analysis to focal regions of earliest neuropathology in Alzheimer’s Disease.

We report evidence that functional connectivity changes occur in the CN to early MCI transition in the absence of structural imaging findings. The progression of observed CN to MCI functional connectivity changes was further evaluated by comparing the mild Alzheimer’s Disease and CN groups. We conclude that rsfMRI functional connectivity analysis of MTL may provide a complementary approach to established neuroimaging and other biomarkers within a model of Alzheimer’s Disease pathophysiology (Jack, 2013) and an Alzheimer’s Disease research framework (Jack, et al., 2018).

## 2. Materials and Methods

Data used in the preparation of this report were obtained from the Alzheimer’s Disease Neuroimaging Initiative (ADNI) database (adni.loni.usc.edu). The ADNI project was launched in 2003 as a public-private partnership, led by Principal Investigator Michael Weiner, MD. The primary goal of ADNI has been to test whether serial magnetic resonance imaging, positron emission tomography, other biological markers, and clinical and neuropsychological assessment can be combined to measure the progression of mild cognitive impairment and early Alzheimer’s Disease.

### 2.1. Participants

The participants in this report were a subset of the ADNI 2 experimental sub-study on resting state fMRI functional connectivity (rsfMRI) (Jack, et al., 2008; Weiner, et al., 2015). The ADNI 2 rsfMRI studies available for download on the Laboratory of Neuro Imaging (LONI) website in September 2016 consisted of 886 scan pairs (structural MRI, rsfMRI) from 220 participants. We subjected scans to a rigorous multistage preprocessing and quality assurance procedure described in further detail below and summarized in Supplemental Figure 1. The procedure yielded a 105 participants cross-sectional study dataset.

Table 1 summarizes participant demographics and metadata. Participants were assigned to one of three clinical categories: CN; early MCI; and mild Alzheimer’s Disease. Clinical assignments adhered to ADNI 2 protocols (publicly available on the adni.loni.usc.edu website). The three clinical groups were balanced for age (*P = 0.56*), education (*P = 0.95*) and gender (*P = 0.22*). The effect of clinical group was significant for Clinical Dementia Rating (CDR) (*P < 0.001*), Clinical Dementia Rating Sum-of-Boxes (CDR-SB) (*P < 0.001*), Mini-Mental State Examination (MMSE) (*P < 0.001*), and APOEε4 (*P = 0.007*).

**Table 1.** Study participant demographics and metadata.

|  | **CN**<br>**n = 32** | **MCI**<br>**n = 39** | **AD**<br>**n = 34** | **P value** |
| --- | --- | --- | --- | --- |
| Age (Mean, SD) | 74.2 (5.5) | 72.5 (6.8) | 73.4 (7.2) | 0.56<sup>(b)</sup> |
| Female (n, %) | 18 (56) | 22 (56) | 13 (38) | 0.22<sup>(a)</sup> |
| Education (Mean, SD) | 16 (2.2) | 16 (2.5) | 17 (2.6) | 0.95<sup>(b)</sup> |
| CDR (Median, Min, Max) | 0 (0, 0) | 0.5 (0.5, 0.5) | 0.5 (0.5, 1.0) | <0.001<sup>(b)</sup> |
| CDR-SB (Median, Min, Max) | 0 (0, 0) | 1.0 (0.5, 3.0) | 2.5 (0.5, 7.0) | <0.001<sup>(b)</sup> |
| MMSE (Median, Min, Max) | 29.5 (27, 30) | 29 (27, 30) | 25 (20, 26) | <0.001<sup>(b)</sup> |
| APOEε4=2 (n, %) | 1 (3) | 4 (10) | 9 (26) | 0.007<sup>(a)</sup> |
| <sup>a</sup>(Chi-square test); <sup>b</sup>(ANOVA);<br>AD = Alzheimer's Disease; CDR = clinical dementia rating; CDR-SB = clinical dementia rating sum-of-boxes,<br>CN = cognitively normal; MCI = mild cognitive impairment; MMSE = mini-mental state examination; SD =<br>standard deviation; |  |  |  |  |

### 2.2. Image acquisition

The present analysis focuses on two of the up to eight scans obtained during a typical ADNI 2 subject visit. These are the high-resolution T_1_-weighted 3D whole brain structural scan (Scan #2 in the protocol) and the resting state functional scan (Scan #4 in the protocol). Structural and rsfMRI scans were acquired with 3T MRI scanners of a single manufacturer (Philips Medical Systems: Achieva, Ingenia, Ingenuity, Intera) from 14 unique sites in two configurations (MULTI-COIL; SENSE-HEAD), and fifteen (15) different software versions (3.2.1 or higher). Structural scans were acquired with a T_1_-weighted sagittal magnetization-prepared rapid gradient echo (MP-RAGE) sequence, with 1.2 mm right/left, 1.0 mm anterior/posterior and 1.0 mm superior/inferior resolution. RsfMRI scans were acquired with a T_2_-weighted gradient echo planar imaging sequence with repetition time/echo time, 3000/30, flip angle, 80°, 48 axial slices, and 64 x 64 in-plane matrix yielding an isotropic 3.3 mm voxel size. Phase-directions varied: anterior-posterior, N = 16; posterior-anterior, N = 86; missing values = 4. Only two temporal slice order sequences were included in this study dataset: a) ascending 48 planes (odd, even), N=98; and b) descending 48 planes (even, odd), N=7. RsfMRI scans were of two durations: a) 140 TRs (“resting state”, N=91); and b) 200 TRs (“extended resting state”, N=14). Additional image acquisition protocol details can be found on the public adni.loni.usc.edu website.

### 2.3. Image processing

Structural images and rsfMRI images were downloaded from LONI either in DICOM format (and locally converted using dcm2nii) or in 4DNIfTI format (DICOM to NIfTI conversion performed by LONI). Preprocessing and data analysis were performed using a combination of FreeSurfer (Dale, et al., 1999), Analysis of Functional Neuroimages (AFNI) (Cox, 1996), and “home-grown” analysis implemented in R and *tcsh* shell scripts. The AFNI Version was AFNI_17.1.03 (2-May-2017). The FreeSurfer version was V6.0 run in the Amazon Cloud on Ubuntu 14.04. The R version was Version 3.4.3 (30-Nov-2017). The AFNI Python script uber_subject.py was used to generate a preprocessing shell script which was manually edited.

#### Structural image analysis with FreeSurfer 6.0

Subcortical and hippocampal subfield segmentation, and cortical parcellation (surface area, thickness and volume) were obtained with FreeSurfer Version 6.0 image analysis suite, which is documented and freely available for download online (http://surfer.nmr.mgh.harvard.edu/) (Dale, et al., 1999). FreeSurfer default settings were applied via the “recon-all” command.

#### Functional MRI analysis with AFNI

The rsfMRI echo-planar image (EPI) preprocessing sequence consisted of the following steps using AFNI: a) alignment of EPI centers to a standard atlas (MNI_avg152T1); b) elimination of the first ten (10) EPIs; c) outlier detection and de-spiking; d) application of time-shift correction; e) alignment to the least motion distorted volume; f) alignment to the anatomical image and warp to a standard space (MNI_avg152T1); g) generation of anatomical and EPI masks; h) application of spatial smoothing (FWHM = 4.0mm); i) scaling voxel time series to a mean of 100; j) generation of demeaned motion parameters and motion parameter derivatives; k) generation of motion “censor” masks (motion limit = 0.3); l) generation of ventricle (Vent) and white-matter (We) segmentation masks from FreeSurfer segmentation and parcellation results; and m) deconvolution with 12 regressors (6 regressors + 6 motion derivatives). The process yielded a 4D residual error volume that served as input to the next stage of volumetric and correlation analysis. Global signal regression (GSR) was not performed. Bandpass filtering was performed in the final ROI pairwise analysis stage as discussed below.

Table 2 lists pre-processing stage group-level statistical summaries obtained from AFNI *@ss_review_basic*. There were no significant differences between groups for average censored motion, maximum censored motion and the average fraction of TR frames censored for excess motion. Group average temporal signal-to-noise values were in excess of 170 with standard deviation in the range 16-19, indicating good overall input EPI signal levels. The values did not differ significantly across groups. Finally, the group average Dice coefficient, which measures the extent of overlap between the anatomical and EPI brain masks, was in excess of 0.88 with standard deviation in the range, 0.012-0.015, indicating good overall anatomical – EPI alignment. The values did not differ significantly across groups.

**Table 2.** Group-average pre-processing statistics.

|  | **CN**<br>**n = 32** | **MCI**<br>**n = 39** | **AD**<br>**n = 34** | **P value<sup>(a)</sup>** |
| --- | --- | --- | --- | --- |
| Avg. Censored Motion (Mean, SD) | 0.095 (0.02) | 0.091 (0.02) | 0.091 (0.02) | 0.63 |
| Max. Censored Motion (Mean, SD) | 0.473 (0.11) | 0.508 (0.13) | 0.506 (0.14) | 0.46 |
| Censor Fraction (Mean, SD) | 0.09 (0.06) | 0.08 (0.06) | 0.10 (0.07) | 0.56 |
| Avg. TSNR (Mean, SD) | 179 (16) | 178 (19) | 180 (16) | 0.85 |
| Global Correlation (Mean, SD) | 0.022 (0.01) | 0.022 (0.01)) | 0.023 (0.01) | 0.91 |
| Dice Coefficient (Mean, SD) | 0.888 (0.01) | 0.891 (0.02) | 0.890 (0.01) | 0.69 |
| <sup>a</sup>(ANOVA); AD = Alzheimer's Disease; CDR = clinical dementia rating; CDR-SB = clinical dementia rating sum-of-boxes, CN = cognitively normal; MCI = mild cognitive impairment; SD = standard deviation; TSNR = temporal signal-to-noise. |  |  |  |  |

### 2.4. Image quality assurance

The initial database consisted of 886 rsfMRI scans (and their paired structural images) from 220 subjects. An R package “ADNIMERGE” contained MRI and rsfMRI quality assessments (e.g., comments by ADNI quality reviewers) and related data (e.g., rsfMRI slice order) stored in the tables “mayoadirl_mri_imageqc” and “mayoadirl_mri_fmri”, respectively. The 886-scan set was reduced to 673 scans (n=196) by rejecting all scans that did not conform to the slice order sequences listed above, or failed visual inspection in the FSL viewer, or contained any reviewer comment in either of the available pair of comment fields. Comments typically referenced motion or clinical findings.

The rsfMRI EPI preprocessing sequence, with motion-limit set to 0.3, was applied to each of the 673 structural-rsfMRI pairs. The global signal was analyzed as an initial screen for excessive motion. Scans were rejected if the global average time-series amplitude value fell outside the range 96-104 or if the standard deviation of the global average time series exceeded 2.0. This gross motion screening procedure reduced the analysis database to 507 structural-rsfMRI pairs (n=190 subjects).

The 507 scan pairs database was further screened using statistics generated by the AFNI-based preprocessing pipeline. Scans were marked for rejection according to the following criteria: a) DICE coefficient < 0.78; b) censor fraction > 0.21; c) maximum censored displacement > 1.00; d) average TSNR > 400 or average TSNR < 100; e) degrees of freedom left < 100; and f) global correlation < 0.10. 206 scan pairs from 106 subjects met these quality assurance criteria. Where multiple candidate scan pairs were present for a given subject, the latest available pair was selected. 80% of the final dataset consisted of scans obtained from subjects who were on at least their second ADNI 2 study visit. The final study database consisted of 105 structural-functional scan pairs from 105 subjects. ANOVA confirmed that preprocessing quality measures were matchead across the three diagnostic groups. See Table 2.

### 2.5. ROI seed and ROI pairs of interest identification

ROI seeds for ROI pairwise functional connectivity analysis were identified by an hybrid *a priori data-driven* method. The procedure was applied independently in left and right hemispheres. To bootstrap the ROI seed selection procedure, FreeSurfer subcortical segmentations were used to define the global 3D bounding box for the hippocampus and MTL. A grid of 6mm-spaced coordinates within the bounding box formed the input candidate ROI seed pool. Input candidate seeds were retained subject to the conditions that the mask for a 6mm radius spatial sample (33 voxels) centered on the ROI seed was 100% contained within the global anatomical brain mask, and not greater than 50% overlap with either of the global ventricle or global white matter masks obtained during pre-processing. The qualifying set of 6mm radius input ROI masks were summed (AFNI *3dMask*) and visually inspected (Supplemental Figure 2). For each subject and for each input ROI mask, the average time series was computed and retained using AFNI *3dmaskave*.

To obtain candidate ROI pairs for analysis, we performed for each input candidate ROI the following group-level procedure. First, the average time series computed and stored in the preceding step, was used to compute the Fisher z-transformed Pearson correlation coefficient with every other voxel in the subject 4D image (AFNI *3dfim+*). Computation was restricted to those voxels contained within the global brain mask generated during AFNI baseline processing). Second, a group-level, one-way, three-factor (CN, MCI, Dementia) 3D ANOVA (AFNI *3dAnova*) was conducted and the 3D volume of F statistics was retained. Third, a 3D cluster analysis (AFNI *3dclust*) was performed on the F statistic 3D volume and for each qualifying cluster a candidate ROI mask of a 6mm spatial sample located at the cluster center of mass was retained. Qualifying clusters were those with threshold F-statistic (F ≥ 3.086; p ≤ 0.05, uncorrected) and minimum cluster size (N_C_ = 20). These parameters were permissive by design. Fourth, only those candidate ROIs were retained for which the mask was 100% contained within the global brain mask (not including the brainstem and cerebellum) and did not have greater than 50% overlap with either the global ventricle or global white matter masks. Fifth, qualifying candidate ROI masks were visually inspected. Sixth, the ROI seed list was pruned to retain only those MTL ROI seeds (the anchor ROIs in ROI pairs of interest) that generated candidate ROIs through the F-statistical map analysis. Finally, individual subject average time series for the final F-statistic map-driven ROI masks were computed and stored.

The above ROI identification procedure yielded ROI “pairs of interest” in which the anchor element in each ROI pair was a left (right) hemisphere MTL ROI. The second element (non-anchor) in each ROI pair was the corresponding F-statistic map-driven ROI whose position was determined by the data. For example, in an ROI pair of interest whose anchor element was in left MTL, the position of the second ROI could lie in ipsilateral or contralateral hemisphere, including MTL. For the left hemisphere, 22 MTL ROI seeds formed the anchor MTL element in 50 ROI pairs of interest (see Supplemental Figure 3). For the right hemisphere, 38 MTL ROI seeds formed the anchor MTL element in 82 ROI pairs of interest (see Supplemental Figure 4). A cluster analysis (AFNI *3dclust*) of the sum of non-anchor ROI masks showed that they comprised a restricted subset of brain regions. For left hemisphere, there were 31 clusters with a maximum of three ROI masks in overlapping anatomical positions. For right hemisphere, there were also 31 clusters with a maximum of five ROI masks in overlapping anatomical positions. Findings in the present study described in detail below are with respect to these anchor MTL and non-anchor ROI pairs of interest.

### 2.6. ROI time series quality assurance

Individual ROI residual time series were band-pass filtered (4^th^ order Butterworth, *f* = {0.01, 0.1}) and averaged, and a subjected to a final quality assurance check for outliers. The 3^rd^-130^th^ (of the original 140 or 200 TRs) time series elements were retained as the candidate final ROI time series. A subject ROI time series was excluded from group-level analysis if any of the following conditions were true: a) the time series was constant; b) the time series standard deviation, σ > 20.0; or c) the fraction of time points with values greater than the outlier threshold θ exceeded 5%. The outlier threshold was set as θ = µ + 3σ, where µ was the time series average value.

### 2.7. ROI pairwise correlation analysis

ROI pairs were subjected to group-level correlation analysis and ANOVA. The core statistical method was one-way, three-factor (clinical group) ANOVA, with *post hoc t* tests applied pairwise (between factors) only for significant ANOVA tests. Correction for large-scale multiple testing was implemented through False Detection Rate (FDR) correction (Benjamini & Hochberg, 1995) and permutation repeated measures (N = 5000) (Nichols and Holmes, 2002; Efron, 2010; Eklund, et al., 2016).

### 2.8. Assignment of anatomical position to an ROI coordinate

ROI seed coordinates were mapped to anatomical locations by analyzing the output of the AFNI *whereami* output. The AFNI *whereami* tool maps an input coordinate to anatomical position with respect to a multiplicity of atlases. To assign an anatomical label we used the “Focus Point”, and “Within 2mm” or “Within 4mm” outputs from the Talairach-Tournoux, MNI (Macro Labels, N27), and MNI (Cytoarchitectonic Max Probability Maps) and checked for consistency. Results were visually confirmed.

## 3. Results

### 3.1. Cortical thinning in group-level AD-CN and AD-MCI contrasts, but not in the MCI-CN contrast

The effect of clinical category on cortical thinning was evaluated using one-way, three-factor ANOVA on estimates of cortical thickness from 34 cortical regions per hemisphere (68 total) (Supplemental Table 1; Supplemental Figure 5). An FDR corrected (*q=0.1, N=68 tests*) ANOVA F statistic critical *p* value threshold, 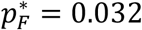, identified 23 significant tests. For significant ANOVA tests only, *post hoc* pairwise *t* tests were computed and FDR-corrected (*q=0.1, N=23 tests*) per contrast and gave critical *p* value thresholds: MCI-CN 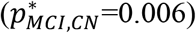; AD-CN 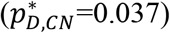; and AD-MCI 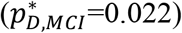. There were twenty-one significant AD-MCI comparisons and fifteen significant AD-CN comparisons. Effect sizes, measured by Cohen’s *d*, were in the range 0.6 ≤ *d* ≤ 1.0. There were two significant positive MCI-CN comparisons (left and right paracentral lobule). All AD-MCI and AD-CN contrasts were negative. There were no significant negative MCI-CN comparisons.

### 3.2. Subcortical volume changes in group-level AD-CN and AD-MCI contrasts, but not in the MCI CN contrast

The effect of clinical category on subcortical volumes was evaluated using one-way, three-factor ANOVA on estimates of subcortical volumes from 8 cortical regions per hemisphere (16 total) (Supplemental Table 2; Supplemental Figure 5). An FDR-corrected (*q*=0.1, *N=16 tests*) oneway three-factor ANOVA F statistic *p* value, 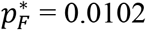, identified eight significant tests. For the significant ANOVA tests only, *post hoc* pairwise *t* tests were computed and FDR-corrected (*q*=0.1, *N=8 tests*) per contrast: AD-CN 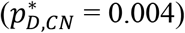; and AD-MCI 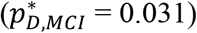. There were eight significant AD-CN contrasts; and six significant AD-MCI contrasts. Cohen’s *d* values were in the range 0.3 ≤ *d* ≤ 1.1. All significant pairwise comparisons for ventricles (e.g., inferior lateral ventricle) were positive and all significant pairwise contrasts for subcortical structures (e.g., whole hippocampus) were negative. With an FDR rate *q* = 0.1, by design set to be liberal, no *p* value fell below the FDR-corrected value (*N=8 tests*) in the MCI-CN contrast, which indicated no significant MCI-CN comparisons.

### 3.3. Hippocampal subfield volume reduction in group-level AD-CN and AD-MCI contrasts, but not the MCI-CN contrast

Hippocampal subfields (11 per hemisphere) were analyzed similarly as above and all twenty-two ANOVA tests were significant, FDR corrected p value (*q=0.1, N=22 tests*), 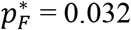 (Supplemental Table 3). *Post hoc* pairwise *t* tests detected twenty-two significant AD-CN comparisons 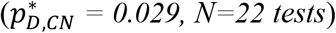, and eighteen significant AD-MCI comparisons (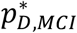 *= 0.077, N=22 tests)*. Effect sizes, measured as Cohen’s *d*, were at least “strong” and above, with left hemisphere effect scores uniformly higher than the right hemisphere (left: 0.9 ≤ *d* ≤ 1.1; right: 0.7 ≤ *d* ≤ 0.86). All significant AD-MCI and AD-CN hippocampal sub-field pairwise contrasts were negative. For the MCI-CN contrast no *p* value was below the FDR-corrected value (*q=0.1, N=22 tests*).

### 3.4. Significant pairwise ROI functional connectivity changes in the MCI-CN group-level contrast

The effect of clinical category on group average ROI pairwise functional connectivity was evaluated using one-way, three-factor ANOVA in separate analyses for the left hemisphere (50 ROI pairs of interest comprising 50 hypotheses, or simultaneous tests) and for the right hemisphere (82 ROI pairs of interest comprising 82 simultaneous tests). For the left hemisphere, an FDR-corrected (*q*=0.05, *N=50 tests*) ANOVA F statistic critical *p* value threshold, 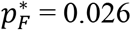, identified 28 significant tests (Supplemental Table 4). For the significant ANOVA tests only, *post hoc* pairwise *t* tests were computed and FDR-corrected (*q*=0.05, *N=28 tests*) independently per contrast: MCI-CN 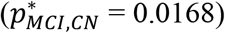; AD-CN 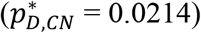; and AD-MCI 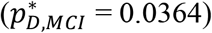. Of the 28 significant ANOVA comparisons, there were ten significant *post hoc* MCI-CN *t* test comparisons (*post hoc* pairwise *t* test *p* < 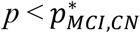). Results are summarized in Table 3. Six of the ten MCI-CN comparisons were positive (indicating hyperconnectivity) with Cohen’s effect size *d*, 0.52 ≤ *d* ≤ 0.92. Four of the ten were negative (indicating hypoconnectivity) with Cohen’s *d*, 0.52 ≤ *d* ≤ 0.79.

**Table 3.** Left MTL ROI - ROI pairwise significant MCI - CN comparisons^4^.

| Anatomical Region(s) |  | ANOVA (1-way, 3-levels) |  |  | Group Average Fisher Z Value |  |  | MCI-CN |  | AD-CN |  | AD-MCI |  |
| --- | --- | --- | --- | --- | --- | --- | --- | --- | --- | --- | --- | --- | --- |
| | | FDR $p^* = 0.026$, $q = 0.05$, $n = 50$ | | | | | | FDR $p^* = 0.0168$ | | FDR $p^* = 0.0214$ | | FDR $p^* = 0.0364$ | |
| MTL ROI | ROI | F | $p$ | $p^*$ | CN | MCI | AD | $p$ | $d$ | $p$ | $d$ | $p$ | $d$ |
| ROI pair hyper-connectivity MCI vs CN |  |  |  |  |  |  |  |  |  |  |  |  |  |
| L: Uncus, A36, Hipp (EC) | L: A38 | 11.3* | 0.00004 | 0.0000 | -0.03 | 0.18 | 0.24 | **<0.001** | **0.92** | **<0.001** | **1.09** | 0.1488 | 0.25 |
| L: Hipp (CA, FD) | R: A38 | 8.7 | 0.00034 | 0.0010 | 0.05 | 0.13 | 0.23 | **0.0156** | **0.52** | **<0.001** | **1.01** | **0.0138** | **0.52** |
| L: A35, A36, Hipp (SUB, CA) | R: Uncus, A28, A36 | 7.0* | 0.00145 | 0.0018 | -0.04 | 0.14 | 0.04 | **<0.001** | **0.83** | 0.0406 | 0.42 | **0.0162** | **0.51** |
| L: Hipp (CA) | R: A10, A47 | 6.3 | 0.00252 | 0.0032 | -0.08 | 0.05 | -0.08 | **0.0016** | **0.66** | 0.4716 | 0.02 | **0.0006** | **0.80** |
| L: A27, Hipp (FD, CA, SUB) | R: A10, A11 | 5.6 | 0.00489 | 0.0056 | 0.00 | 0.08 | -0.04 | **0.0168** | **0.52** | 0.1380 | 0.27 | **0.0012** | **0.74** |
| L: Uncus, A36, A28, Hipp (SUB) | L: A34 | 4.4 | 0.01515 | 0.0188 | -0.06 | 0.08 | 0.06 | **0.0042** | **0.65** | **0.0096** | **0.57** | 0.3524 | 0.09 |
| ROI pair hypo-connectivity MCI vs CN |  |  |  |  |  |  |  |  |  |  |  |  |  |
| L: Hipp (FD, CA, SUB), Ling. G. | L+R: Precuneus | 5.8 | 0.00424 | 0.0040 | 0.20 | 0.05 | 0.09 | **0.0002** | **0.79** | **0.0050** | **0.63** | 0.2098 | 0.19 |
| L: A35, A36, Hipp (SUB, CA) | L+R: Precuneus | 4.6 | 0.01187 | 0.0122 | 0.21 | 0.08 | 0.10 | **0.0034** | **0.68** | **0.0142** | **0.56** | 0.3586 | 0.09 |
| L: A28, A35, Hipp (SUB, EC, CA) | R: A45, A47 | 4.5 | 0.01291 | 0.0104 | 0.08 | -0.05 | 0.07 | **0.0098** | **0.58** | 0.4490 | 0.04 | **0.0040** | **0.64** |
| L: Caudate tail, Hipp (CA, FD) | L+R: ACC | 3.8 | 0.02570 | 0.0260 | 0.12 | 0.01 | 0.06 | **0.0016** | **0.71** | 0.0622 | 0.38 | 0.1258 | 0.27 |
| ACC = anterior cingulate cortex; AD = Alzheimer's disease; CA = cornu ammonis; CN = cognitively normal; EC = entorhinal cortex; FD = dentate gyrus; FDR = false detection rate; Hipp = hippocampus; Ling. G. = lingual gyrus; MCI = mild cognitive impairment; MTL = medial temporal lobe; ROI = region of interest; SUB = subiculum; *Significant at FDR $q = 0.01$. | | | | | | | | | | | | | |

ACC = anterior cingulate cortex; AD = Alzheimer's disease; CA = cornu ammonis; CN = cognitively normal; EC = entorhinal cortex; FD = dentate gyrus; FDR = false detection rate; Hipp = hippocampus; Ling. G. = lingual gyrus; MCI = mild cognitive impairment; MTL = medial temporal lobe; ROI = region of interest; SUB = subiculum; \* (Significant at FDR $q = 0.01$ ).

For the right hemisphere, an FDR-corrected (*q*=0.05, *N=82 tests*) ANOVA F statistic critical *p* value threshold, 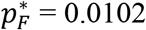, identified 17 significant MCI-CN comparisons (Supplemental Table 5). For the significant ANOVA tests only, *post hoc* pairwise *t* tests were computed and FDR-corrected (*q*=0.05, *N=17 tests*) independently per contrast: MCI-CN 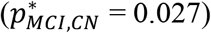; AD-CN 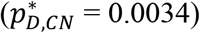 and AD-MCI 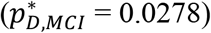. Of the 17 significant ANOVA comparisons, there were 13 significant *post hoc* MCI-CN *t* test comparisons (*post hoc* pairwise *t* test *p* < 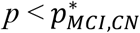). Results are summarized in Table 4. Nine of the 13 MCI-CN comparisons were positive (indicating hyperconnectivity) with Cohen’s effect size *d*, 0.45 ≤ *d* ≤ 0.95. Four of the 13 were negative (indicating hypoconnectivity) with Cohen’s *d*, 0.57 ≤ *d* ≤ 0.73.

**Table 4.**
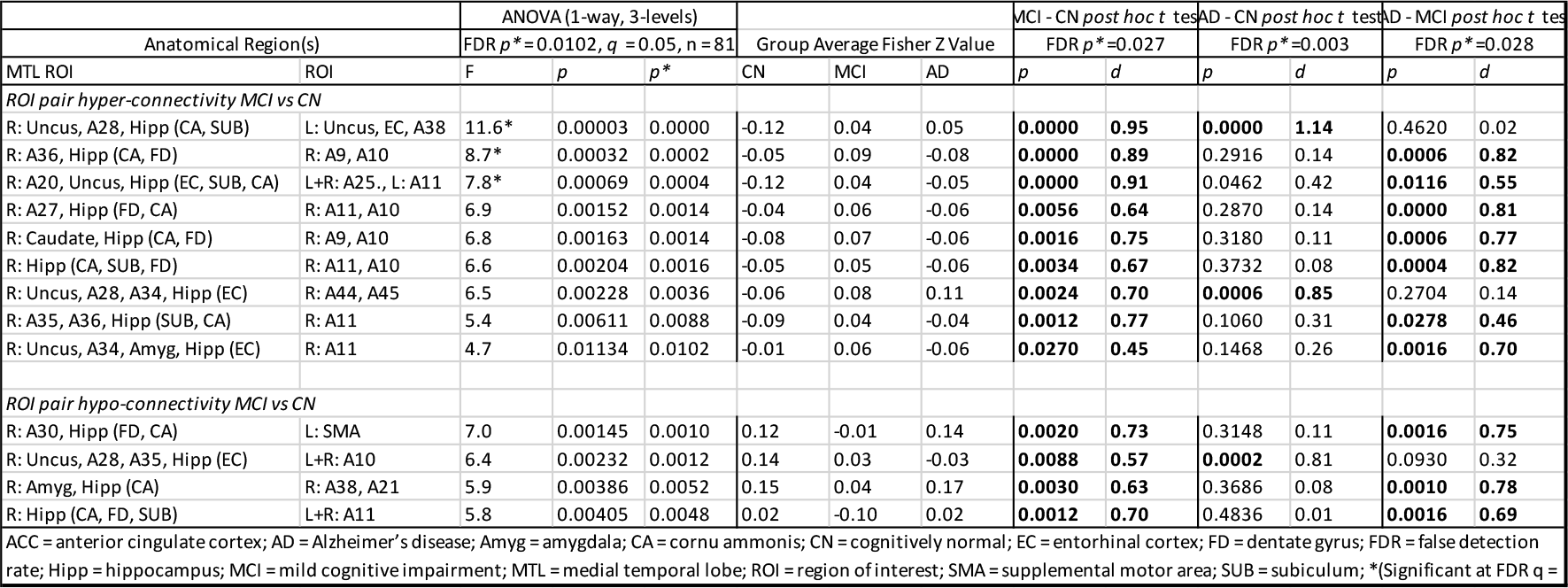
Right MTL ROI - ROI pairwise significant MCI - CN comparisons^5^.

### 3.5. Parcellation of functional connectivity changes of left MTL in the group-level MCI-CN contrast

The positions of left MTL anchor ROI masks for significant ROI pairs of interest in the MCI-CN contrast are shown in sagittal section (Figure 1 (top)) and coronal section (Figure 3 (left)). Positions of the corresponding non-anchor paired ROIs are discussed next and shown in Supplemental Figure 6. The spatial organization of the ten MTL anchor ROI masks was analyzed using AFNI *3dclust* which identified three clusters oriented along a ventral-medial to dorsal-lateral axis in the left MTL. First, in the ventral medial cluster, in the area of the hippocampal head, the most medial anchor ROIs were positioned in the parahippocampal gyrus (uncus), and Areas 35 and 36, and were ipsilaterally hyperconnected with a more rostral non-anchor ROI positioned in Area 34. More laterally in this cluster, anchor ROI positions progressively shifted to Area 28, hippocampus (subiculum and CA) and parahippocampal gyrus. Functional connectivity changes with paired non-anchor ROIs consisted of hyperconnectivity with ROIs in ipsilateral and contralateral Area 38, and hypoconnectivity with a contralateral ROI positioned laterally in Area 45 and 47.

**Figure 1.**
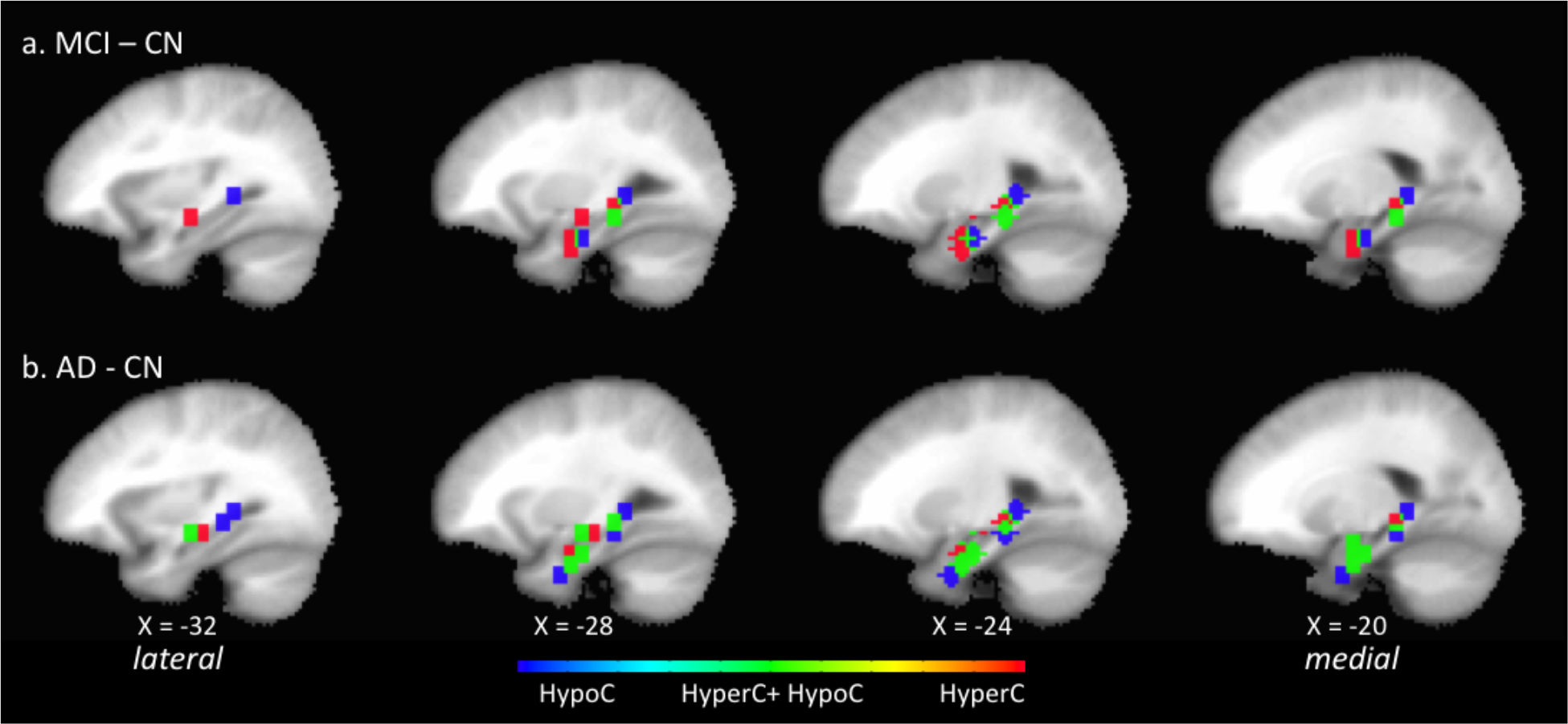
Position in the left hemisphere sagittal plane (4mm stride) of anchor MTL ROI elements of significant ROI pairs of interest in the MCI – CN contrast (top) and D – CN contrast (bottom). Shown are ROI masks for the 6-mm radius volume centered at the ROI seed. In both contrasts, ROIs group in a ventral anterior subzone and a dorsal posterior subzone. There is progressive change that may correlate with disease progression. In the D – CN contrast, there are more significant ROI pairs of interest collectively whose anchor MTL ROIs cover a larger extent of the MTL compared to MCI – CN. In addition, there is an increase in the number of individual MTL ROIs that form multiple significant ROI pairs in which one pair is of type hypoC and the other hyperC. Positions of the ROI paired with each MTL ROI shown are summarized in Table 3, their MNI coordinates are listed in Supplemental Tables 4 and 6, and corresponding image maps are shown Supplemental Figures 6 and 7. The direction of ROI pair functional change is color-coded for hyperconnectivity (*red*), hypoconnectivity (*blue*), and simultaneous hyperconnectivity and hypoconnectivity (*green*).

A second cluster of MTL anchor ROI masks, lateral and dorsal to the ventral medial cluster, was positioned in its medial subregion in the hippocampus (subiculum, CA), and in its lateral subregion in the hippocampus (CA) in the region of the caudate tail and parahippocampal gyrus. The anchor ROI in this cluster was hyperconnected with its paired non-anchor ROI positioned in contralateral lateral Area 10 adjacent to Area 47.

Last, in the dorsal cluster of anchor ROI masks, its medial subregion was positioned in Area 36 and hippocampus (CA, FD, subiculum), and in its lateral subregion in hippocampus (CA) in the region of the caudate tail and parahippocampal gyrus. Functional connectivity changes in this subzone included: a) an anchor ROI hyperconnected with a contralateral non-anchor ROI positioned in lateral Area 10/11; b) an anchor ROI hypoconnected with a non-anchor ROI located in precuneus; c) an anchor ROI hypoconnected with a non-anchor ROI located in anterior cingulate cortex; and d) an anchor ROI simultaneously hyperconnected with a contralateral non-anchor ROI located in the region of uncus, Area 28 and 36, and hypoconnected with a non-anchor ROI located in precuneus.

In summary, in the left hemisphere, for the identified significant ROI pairs in the MCI-CN contrast, the anchor MTL ROIs clustered within focal subregions of entorhinal and perirhinal areas, hippocampus, and parahippocampal gyrus. The spatial pattern of functional connectivity changes with corresponding non-anchor ROIs suggested that MTL subzones were: a) ventrally, bilaterally hyperconnected, including uncus and Area 38; b) dorsally and laterally, simultaneously hyperconnected and hypoconnected with distinct non-overlapping contralateral lateral frontal cortical regions; and c) over an extended dorsal and lateral zone, hypoconnected with midline regions associated with anterior and posterior default-mode network (DMN) nodes.

### 3.6. Parcellation of functional connectivity changes of right MTL in the group-level MCI-CN contrast

The position of right MTL anchor ROI masks for significant ROI pairs of interest in the MCI-CN contrast are shown in sagittal section (Figure 2 (top)) and coronal section (Figure 3 (left)). Positions of the corresponding non-anchor paired ROIs are discussed next and shown in Supplemental Figure 7. The spatial organization of the 13 MTL anchor ROI masks was analyzed using AFNI *3dclust* which identified two clusters oriented along a ventral-medial to dorsal-lateral axis. In the first cluster, near the head of the hippocampus, in its medial subregion, anchor ROIs were positioned in the uncus, and Areas 28, 34, 35 and 36. Anchor ROI positions in the middle and lateral subregions progressively shifted to the hippocampus (subiculum, FD and CA). Pairwise functional connectivity with corresponding non-anchor ROIs consisted of an interleaved pattern of hyperconnectivity and hypoconnectivity with small regions of overlap. In the medial subregion, immediately adjacent anchor ROIs were either hyperconnected with an ipsilateral lateral non-anchor ROI in Area 44 and 45, or hypoconnected bilaterally with a non-anchor ROI in Area 10 at the midline. In the middle subregion there was a zone of hypoconnectivity with an ipsilateral non-anchor ROI in the region of Area 38 and 21. Immediately lateral to this hypoconnected anchor ROI, were a pair of anchor ROIs that were hyperconnected with non-anchor ROIs positioned in contralateral uncus, Area 38 and 28. In the most lateral subregion, anchor ROIs were hyperconnected bilaterally with a non-anchor ROI in the region of Area 11 and 25 at the midline.

**Figure 2.**
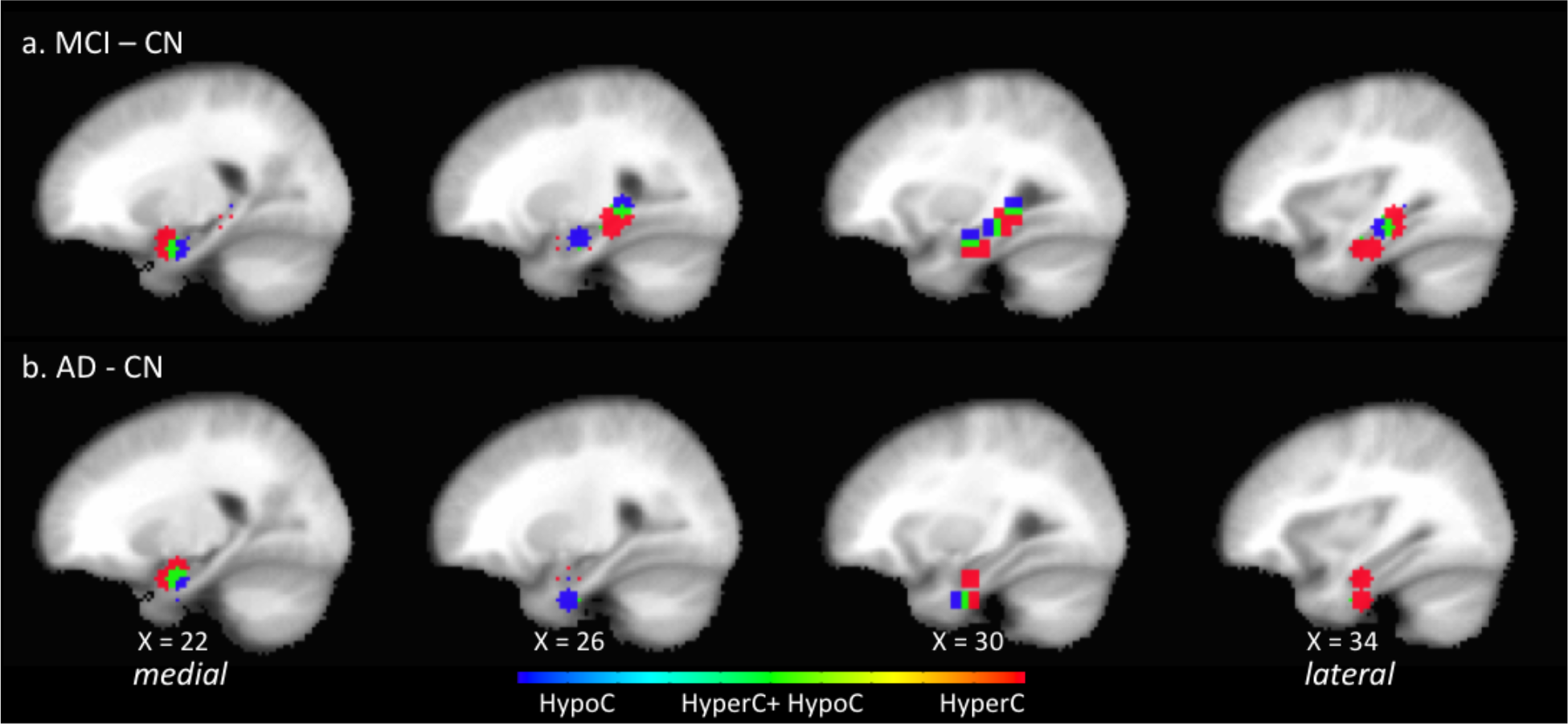
Position in the right hemisphere sagittal plane (4mm stride) of anchor MTL ROI elements of significant ROI pairs of interest in the MCI – CN contrast (top) and D – CN contrast (bottom). Shown are ROI masks for the 6-mm radius volume centered at the ROI seed. In the MCI - CN contrasts, but not the D – CN, ROIs group in a ventral anterior subzone and a dorsal posterior subzone. In the D – CN contrast, the dorsal posterior ROI cluster is not present, indicating that there may be a class of ROI pairwise functional connectivity changes that persist and those that are associated specifically with the CN to MCI transition. Positions of the ROI paired with each MTL ROI shown are summarized in Table 4, their MNI coordinates are listed in Supplemental Tables 5 and 6, and corresponding image maps are shown Supplemental Figures 8 and 9. The direction of ROI pair functional change is color-coded for hyperconnectivity (*red*), hypoconnectivity (*blue*), and simultaneous hyperconnectivity and hypoconnectivity (*green*).

**Figure 3.**
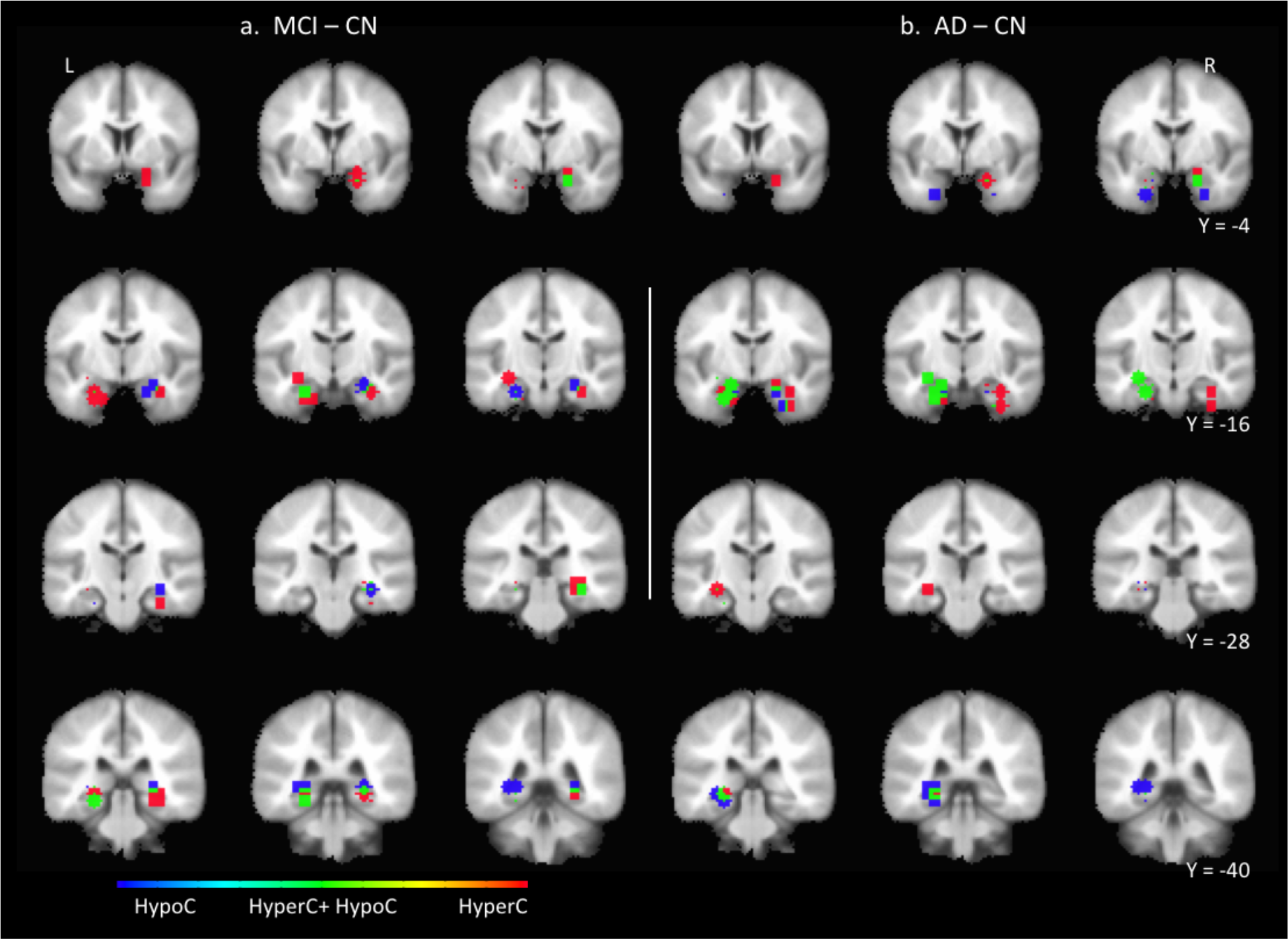
Coronal series showing the position of anchor MTL ROI elements of significant MCI – CN contrast ROI pairs (a) and D – CN contrast (b) ROI pairs. Shown are the anchor MTL ROI masks consisting of a 6-mm radius volume centered at the anchor MTL ROI seed. The direction of ROI pair functional change is color-coded for hyperconnectivity (*red*), hypoconnectivity (*blue*), and simultaneous hyperconnectivity and hypoconnectivity (*green*).

The second cluster of anchor ROIs, dorsal to the first, was characterized by multiple immediately adjacent anchor ROIs with a distinct distribution of functional connectivity with corresponding non-anchor ROIs. In its medial extent, anchor ROIs in this cluster were positioned in Area 35, 36 and hippocampus (CA, FD, subiculum) and in its lateral extent in hippocampus (CA, FD) in the region of the caudate tail, but not parahippocampal gyrus. In the medial extent of this cluster of anchor ROIs, functional connectivity changes with non-anchor ROIs consisted of hyperconnectivity with an ipsilateral lateral ROI positioned in Area 10 and 11. In the lateral extent of this cluster of anchor ROIs, there was hyperconnectivity with an ipsilateral lateral non-anchor ROI whose position shifted to that of Area 9 and 10. In the medial subzone of this cluster of anchor ROIs, there was functional hypoconnectivity with a corresponding paired non-anchor ROI positioned in contralateral supplemental motor area (SMA) and in its lateral extent hyperconnectivity bilaterally with a non-anchor ROI positioned in Area 11 at the midline.

In summary, in the right hemisphere in the transition from CN to MCI, the spatial pattern of pairwise functional connectivity changes between anchor MTL and non-anchor ROIs suggests that MTL subzones were: a) ventrally, hyperconnected with contralateral uncus and Area 38 and hypoconnected with ipsilateral anterior Area 38; b) hyperconnected with ipsilateral lateral frontal cortex; and c) hypoconnected with frontal cortical zones at the midline in regions adjacent to or associated with anterior DMN midline nodes.

### 3.7. Significant pairwise ROI functional connectivity changes in the AD-CN group-level contrast

In the left hemisphere, an ROI pair was significant for functional connectivity changes in the AD-CN contrast for *post hoc* pairwise *t* test critical *p* value threshold less than the FDR-corrected, 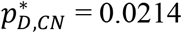, (*q*=0.05, *N=28 tests*). There were 21 significant AD-CN comparisons (Supplemental Tables 4 and 6). The MTL anchor ROI mask positions are shown in Figure 1 (bottom) and Figure 3 (right), and non-anchor ROI mask positions are shown in Supplemental Figure 8. Comprising the 21 significant comparisons were: a) nine hyperconnected pairs, Cohen’s d, 0.52 ≤ *d* ≤ 0.86, median=0.75; and b) 12 hypoconnected pairs, Cohen’s d, 0.7 ≤ *d* ≤ 0.86, median=0.64. There were more than twice as many AD-CN significant comparisons (N=21) as there were MCI-CN significant ROI comparisons (N=10). There were 1.5 times as many hyperconnected ROI pairs (N_AD-CN_=9 vs N_MCI-CN_=6) and 3.0 times as many hypoconnected ROI pairs (N_AD-CN_=12 vs N_MCI-CN_=4). Six of the nine significant hyperconnected and ten of the twelve significant hypoconnected ROI pairs were not significant in the MCI-CN contrast. Despite the increase in numbers of significant ROI pairs, the overall spatial organization of MTL ROI positions was similar to that of the MCI-CN contrast. The greater number of co-located anchor ROIs preferentially distributed in the ventral medial and dorsal lateral ROI clusters.

A striking feature in the AD-CN contrast was individual anchor MTL ROIs which formed multiple significant ROI pairs. Fourteen left hemisphere MTL ROIs comprised the anchor ROI in 21 significant ROI pairs. This was compared to nine MTL anchor ROIs that formed 10 significant ROI pairs in the MCI-CN contrast. In the AD-CN contrast, six MTL ROIs comprised the anchor element in two significant ROI pairs. The most common combination (four of six) was one hyperconnected pair and one hypoconnected pair. For example, an anchor ROI in ventral MTL (Area 35) was hyperconnected with a non-anchor ROI located contralateral Area 38 and simultaneously hypoconnected with a non-anchor ROI in the region of the orbital gyrus at the midline. As a second example, an anchor ROI in the dorsal MTL in hippocampus (CA) was hyperconnected with a contralateral non-anchor ROI located in the region of uncus and entorhinal cortex, and simultaneously hypoconnected with a non-anchor ROI in contralateral lateral inferior temporal gyrus. In the single case of an anchor MTL ROI having two hypoconnections, an anchor ROI positioned in hippocampus (CA) adjacent to the parahippocampal gyrus, was hypoconnected with a non-anchor ROI located in precuneus at the midline and hypoconnected with a non-anchor ROI located in the contralateral lateral frontal cortex in Area 45 immediately adjacent to Area 47. Last, in the single case of an anchor MTL ROI having two hyperconnections, an anchor ROI positioned in the region of Areas 28, 25, 36 and uncus, was *bilaterally* hyperconnected with a non-anchor ROI located in Area 38.

In the right hemisphere, ROI pairwise tests were significant for functional connectivity changes in the AD-CN contrast with a *post hoc* pairwise *t* test critical *p* value threshold less than the FDR-corrected, 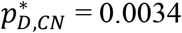, (*q*=0.05, *N=17 tests*). There were six significant comparisons (Supplemental Tables 5 and 6). The MTL anchor ROI mask positions are shown in Figure 2 (bottom) and Figure 3 (right) and the paired non-anchor ROI mask positions are shown in Supplemental Figure 9. Four of the six AD-CN comparisons were positive indicating hyperconnectivity with Cohen’s effect size *d*, 0.7 ≤ *d* ≤ 1.14. Two of these four significant comparisons were also significant in the MCI-CN contrast, indicating persistence of effect with disease progression: hyperconnectivity with contralateral uncus; and hyperconnectivity with ipsilateral Area 44 and 45. Two of these four significant comparisons were not significant in the MCI-CN contrast with the following characteristics: a) both MTL anchor ROIs were positioned in the region of the head of the hippocampus, entorhinal cortex and uncus; b) one (dorsal) was hyperconnected with a contralateral non-anchor ROI located in lateral Area 10; and c) the other (ventral) was hyperconnected with an ipsilateral non-anchor ROI located in medial Area 11 (nearly at the midline). Last, for two of the six significant AD-CN ROI pairs, neither of which was significant in the MCI-CN comparison, the pairwise functional connectivity change was hypoconnectivity (Cohen’s *d*, 0.77 ≤ *d* ≤ 0.81). The anchor ROIs in these pairs were positioned in the region of the head of the hippocampus, entorhinal cortex and uncus. One (dorsal) was hypoconnected with a non-anchor ROI located in Area 10 at the midline; the other (ventral) was hypoconnected with a non-anchor ROI located in ipsilateral lateral A10 adjacent to A46.

Summarizing left and right hemisphere jointly, the spatial pattern of functional connectivity changes from CN to D reflected a combination of consolidation, progression, and reversion of the changes observed in the CN to MCI contrast. First, there were not anchor ROIs positioned in additional MTL subzones. Second, additional significant ROIs in the AD-CN contrast clustered preferentially in the region of the head of the hippocampus, entorhinal cortex and uncus, compared to the wider distribution in the MCI-CN contrast. Third, significant AD-CN non-anchor ROIs were positioned within similar cortical regions (e.g., lateral frontal cortex, midline regions) as in the MCI-CN contrast or in additional regions (e.g., right inferior temporal cortex). Last, a combination of reversion and consolidation was evident in the absence in right hemisphere in the AD-CN contrast of the more dorsal cluster of ROIs present in MCI-CN contrast.

### 3.8. MCI-CN functional connectivity changes may persist or revert in the group-level AD-CN contrast

In the AD-CN contrast there were functional connectivity changes that persisted or were transient compared to the MCI-CN contrast. In the left hemisphere for three of six MCI-CN hyperconnected ROI pairs and for two of four MCI-CN hypoconnected ROI pairs, the functional connectivity change persisted or was amplified in the AD-CN contrast. In the right hemisphere, however, only two of nine MCI-CN hyperconnected ROI pairs and for one of four MCI-CN hypoconnected ROI pairs, did the functional connectivity change persist. Examples of persistent effects included: a) left MTL and its ipsilateral and contralateral hyperconnectivity with Area 38; b) left MTL hypoconnectivity bilaterally with anterior cingulate cortex and precuneus; and c) right MTL hypoconnectivity bilaterally with Area 10 at the midline. Of the set of ROI pairs with persistent functional connectivity changes in the MCI-CN and AD-CN contrasts, only one ROI pair was monotone, i.e., significant in all three contrasts MCI-CN, AD-CN, and AD-MCI. For this progressively hyperconnected ROI pair, the MTL ROI element was positioned in left MTL in the region of Areas 28, 35, 36 and uncus, and the second element was in a contralateral region with focal point in Area 38. In contrast, for three of six hyperconnected and for two of four hypoconnected significant MCI-CN ROI pairs in the left hemisphere, and in all but three of 13 significant ROI pairs in the right hemisphere, the changes reversed course such that there was no significant difference in the AD-CN contrast.

In summary, there were monotonic and non-monotonic, transient and persistent changes in functional connectivity between the identified ROI pairs in the AD-CN contrast compared with the MCI-CN contrast. The enduring changes observed across the CN, MCI and D progression include mutual bilateral hyperconnectivity between MTL and Area 38, and hypoconnectivity of left and right MTL with anterior and posterior regions immediately adjacent to or associated with midline DMN nodes.

### 3.9. Sensitivity analysis

In preliminary studies, we assessed by spot-checking the sensitivity of the detection of statistically significant (one-way, three-factor ANOVA ROI pair) functional connectivity comparisons to changes in preprocessing parameters. The results reported in this study were obtained with FWHM = 4.0 mm spatial smoothing in the AFNI preprocessing stage. Smoothing with FWHM = 6.0 mm or FWHM = 8.0 mm and FDR *q* = 0.05, eliminated the observation of significant ANOVA trials. In such cases, we did not proceed to the 5000 permutation repeated measures test (although we observed anecdotally using a 1000 permutation repeated measures test that a few significant tests were observed). Similarly, present results were obtained using an ROI defined as the 6.0 mm radius volume centered at a given source or target ROI seed. Expansion to an 8.0 mm radius volume eliminated the observation of significant ANOVA tests. In any of the above tests, when we relaxed the FDR *q* value, for example, to 0.05 ≤ *q* ≤ 0.20, progressively greater numbers of significant ANOVA tests were observed. These preliminary observations drove the decision to proceed with FWHM = 4.0 mm and ROI volume radius = 6.0 mm where, with a conservative FDR *q* = 0.05.

We assessed the reproducibility of the reported results under repeated measures with different random number generator seeds in the 5000-trial permuted repeated measures. We completed two additional 5000-trial runs for the left hemisphere data and found that for the MCI-CN contrast (the only contrast analyzed in detail) there were comparable numbers of significant tests and same, adjacent or overlapping ROI pairs with comparable F statistic values to those reported here. The remaining pairs were typically in positions adjacent to or overlapped those reported.

We determined the impact of an even more conservative FDR value *q* = 0.01 on the reported pairwise ROI functional connectivity results. For the left hemisphere, the FDR corrected F statistic *p* value was (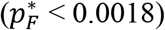) and there remained nine significant ANOVA tests. For the significant ANOVA tests only, *post hoc* pairwise *t* tests were computed and FDR-corrected (*q* = 0.01, N =9 tests) independently per contrast: 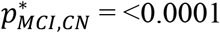; 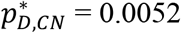; 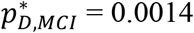. There were no significant hypoconnected ROI pairs. There were two significant hyperconnected MCI-CN pairs and these are shown in Table 3 marked with an asterisk. For the right hemisphere, the FDR corrected F statistic *p* value was (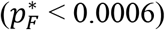) and there remained five significant ANOVA tests. For the significant ANOVA tests only, *post hoc* pairwise *t* tests were computed and FDR-corrected (*q* = 0.01, N =5 tests) independently per contrast: 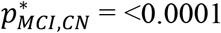; 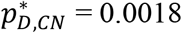; 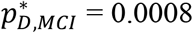. There were no significant hypoconnected ROI pairs. There were three significant hyperconnected MCI-CN pairs and these are shown in Table 4 marked with an asterisk. In summary, the impact of an even more conservative FDR value q = 0.01 on reported MCI-CN significant comparisons is that the findings of hypoconnected ROI pairs are not conserved; the findings of hyperconnected ROI pairs are conserved. For the D-CN contrast, findings are confirmed, although with reduced numbers, in that there remain significant comparisons in left and right hemisphere representing hyperconnected ROI pairs and hypoconnected ROI pairs.

## 4. Discussion

We report evidence that disruption of task-free resting state network functional connectivity is present before the Alzheimer’s Disease pathophysiological stage at which neurodegeneration has sufficiently progressed to be detected. The results were obtained by comparing resting-state fMRI connectivity with an automated analysis of standard (1mm) structural T1 MRI as cortical thinning, subcortical structural volume reduction and hippocampal subfield volume reduction. An anatomically bootstrapped, data-driven procedure identified statistically significant comparisons in resting state functional networks having edges where one node was in the MTL and the other was at anatomical locations corresponding with DMN nodes. Further, we observed a diversity of effects of Alzheimer’s Disease progression on ROI pairwise functional connectivity, including complex patterns of hypoconnectivity *and* hyperconnectivity across the MCI-CN and AD-CN transitions confirming that the effect of Alzheimer’s Disease progression on functional connectivity is more complex than simply there being monotonic loss or gain of functional connectivity.

### 4.1. Structural imaging findings provide an important control for functional connectivity findings

A key structural MRI finding in Alzheimer’s Disease is cortical atrophy (thinning and volume loss) that correlates with symptom severity and has a demonstrated diagnostic and prognostic value (Braak and Braak, 1991; Fan, et al., 2008; Davatizikos, et al., 2009; Lerch, 2005; Jack, et al., 2008; Dickerson, et al., 2009; Misra, et al., 2009; Miller-Thomas, et al., 2016). This finding was reproduced here (Supplemental Table 1) and served as an important control for functional connectivity analysis. We hypothesized that the anatomical locations of the cortical ROIs observed in our functional connectivity analysis would correspond with previously demonstrated spatial patterns of cortical thinning. Indeed, the anatomically boot-strapped data-driven method employed in the present study identified significant MCI-CN cortical ROI anatomical positions in areas known to be most vulnerable to cortical thinning. These included the rostral medial temporal cortex, temporal pole and superior frontal regions, but not the inferior temporal cortex. Significant functional connectivity changes involving inferior temporal cortex were however observed in the AD-CN contrast. The absence of statistically significant cortical thinning in the MCI-CN contrast suggests that the observed functional connectivity changes may not solely be accounted for as a reflection in MTL of mechanisms arising in cortical regions.

The present study also reproduces key findings of hippocampal volume and hippocampal subfield volume loss with Alzheimer’s Disease progression (Hyman, et al., 1984; Braak and Braak, 1991; Jack, et al., 1992, 1998; Schuff, et al., 2009; Boutet, et al., 2014; Sorensen, et al., 2016; Wolk, et al., 2017). The confirmation of expected significant volume loss and significant functional connectivity changes in the AD-CN contrast provides a control for the present MCI-CN and AD-CN findings.

The absence of significant reduction in whole hippocampus volume and hippocampal subfield volumes in the MCI-CN contrast reported above is with respect to a widely available benchmark: up-to-date automated analysis (FreeSurfer) of research quality T1-weighted scans (1.0 x 1.0 x 1.0 mm^3^). We do not conclude that there are no morphometric changes taking place contemporaneously with the CN to MCI transition. Wolk, et al. (2017), analyzed high resolution (~0.4 x 0.4 x 2 mm^3^) T2-weighted MRI scans orthogonal to the long axis of the hippocampus at autopsy. They reported significant group-level reduction of MTL subfield volumes (e.g., Area 35, perirhinal cortex) in the CN to early MCI transition where whole hippocampus volumes were not significantly different. In a feasibility study of an *in vivo* approach, Boutet, et al., (2017) used 7T MRI to detect hippocampal subfield volume changes in an AD–CN contrast. These studies, together with present results, indicate the potential of dual functional and high-resolution morphometric biomarkers for detecting the earliest stages in the Alzheimer’s Disease pathophysiology.

### 4.2. Comparison to prior work

With respect to our functional connectivity findings, five published reports are of particular relevance (Grecius, et al., 2004; Jones, et al, 2011; Brier, et al., 2012; Ward, et al., 2014; Dillen, et al., 2017). Grecius, et al., (2004) performed a pairwise group-level (healthy control (N=14) vs very mild or mild Alzheimer’s Disease (N=13)) independent components analysis (ICA) and ROI analyses of steady-state DMN connectivity during a simple sensorimotor task in a cohort drawn from the Washington University Alzheimer’s Disease Research Center. Key findings were: a) prominent bilateral coactivation of the hippocampus with DMN in both clinical groups; and b) disruption of hippocampal – PCC connectivity in the Alzheimer’s Disease group. Dillen et al., (2017) performed a within-group and between-group analysis of hippocampus and DMN node functional connectivity analysis in a cohort drawn from an outpatient population at the Memory Clinic Cologne Julich (healthy control (N=25), subjective cognitive decline (SCD) (N=28); and prodromal Alzheimer’s Disease (N=25)). Significantly, they reported a decoupling of hippocampus with posterior cortical DMN nodes in both SCD and prodromal Alzheimer’s Disease groups.

Brier, et al., (2012) performed a cross-sectional analysis of five task-free resting state networks (RSN), including DMN, in a cohort drawn from the Knight Alzheimer’s Disease Research Center at Washington University in St. Louis. Subjects were assigned to one of three groups according to CDR scores 0 (cognitively normal, N=386), 0.5 (very mild AD, N=91), and 1.0 (mild AD, N=33). The study reported summary measures of intranetwork and internetwork steady-state correlation structure based on Fisher z-transformed pairwise Pearson correlation analysis of ROI seed-based average time series. While not uniformly statistically significant, the conclusion was that there was group-level progressive loss of intranetwork and internetwork connectivity. The study also reported results of the effect of clinical group (CN, MCI, Alzheimer’s Disease) on ROI pair-wise functional connectivity between PCC and eight other DMN nodes. For two ROI pairs (PCC – medial prefrontal cortex; PCC –left inferior temporal lobe) there was a statistically significant reduction of connectivity in the CDR 0.5 group compared to the CDR 0 group. Present results confirm the Brier, et al. observation of ROI pairwise functional connectivity loss in the MCI-CN comparison. They also extend the Brier, et al. results. First, where Brier, et al., utilized a single ROI seed to estimate midline DMN structures (e.g., PCC, thalamus), the present study utilized an anatomically bootstrapped data-driven method that identified distinct left and right ROI seeds in these structures. The result was a more fine-grained ROI pair-wise analysis which led to the observation of widespread left- and right-hemispheric MTL hyperconnectivity ventrally together with disconnection with midline DMN and DMN coactivated regions. Second, by design, the present study included hippocampus and other MTL structures in the analysis and presented evidence that these structures are involved in the loss of functional connectivity in the MCI-CN comparison.

Jones, et al. (2011) performed group-level functional connectivity analyses of task-free steady-state DMN connectivity in cohorts drawn from the Mayo Clinic Alzheimer’s Disease Research Center and Mayo Clinic Study of Aging. Age-related changes in DMN functional connectivity were analyzed using an ICA approach in a cognitively normal cohort (N=341). The age-related changes consisted of differential effects in anterior and posterior DMN connectivity: a) anterior DMN showed both increases and decreases in connectivity within the frontal lobe; and b) posterior DMN showed a pattern of predominantly but not exclusively decreases in connectivity. An age-matched CN and Alzheimer’s Disease group analysis using both ICA and seed-based ROI approaches concluded that DMN connectivity changes in Alzheimer’s Disease represent an acceleration of the same aging pattern observed in the control sample. The present study complements these findings by providing additional detailed parcellation of changes in MTL, separately analyzed in left and right hemisphere.

Ward, et al., (2014), in their study comparing resting state data and an associative memory encoding task, concluded that the parahippocampal gyrus is the primary hub of the DMN in the MTL during resting state. Their report ends with the suggestion that measuring PHG connectivity may provide a biomarker of early Alzheimer’s Disease. The present study, although motivated by a different hypothesis (that functional connectivity changes temporally precede detection of significant neurodegeneration) and using an entirely independent analysis method (anatomically bootstrapped, data-driven method), provides quantitative evidence in support of the Ward, et al., suggestion. We observe evidence of functional disconnection bilaterally between posterior cortical DMN nodes (anterior cingulate cortex, precuneous) and parahippocampal gyrus as part of a larger pattern of disruption involving anterior and posterior DMN, and subcortical structures including hippocampus and parahippocampal gyrus.

### 4.3. Limitations

Limitations of the present study relate to the study population, the imaging data and preprocessing methods. First, by excluding participants with other neurological injury or findings, the study population was not representative of a general age-matched clinical population. Second, the effect of the “penciling artefact”, found in ADNI image data in left lateral frontal cortex, on functional connectivity measures is unknown and introduces uncertainty in the interpretation of present results. For example, there was a multiplicity of significant ROI pairs between right MTL and ipsilateral lateral frontal cortex. There was not a comparable finding in the left hemisphere, whereas there were bilateral findings with respect to midline ROIs. Third, the present study applied in the pre-processing stage an atlas (AFNI: MNIavg152) derived from younger brains than used in this study which could inject systematic error into the data. Similarly, the FreeSurfer parameters were the default set which may be sub-optimal given the population comprising this study. Last, there is not a consensus regarding the use or non-use of GSR (Murphy and Fox, 2017) and the present study did not apply GSR. The reasoning was two-fold. First, given the “penciling artefact” in ADNI data, it was problematic whether and how to obtain a viable global signal estimate. Second, the preprocessing procedure estimated and removed motion confounds and the ROI identification procedure applied ventricle and white matter masks to minimize the influence of non-grey matter signals. However, any global or large-scale regional neuronal fluctuations would not be controlled in this procedure.

### 4.4. Translational considerations

A challenge in the consideration of functional connectivity biomarkers is that there are in clinical use well-established semi-quantitative structural biomarkers of Alzheimer’s Disease (Fazekas, et al., 1987; Scheltens, et al., 1992; Wahlund, et al., 2001). These methods have been extended in clinical practice and medical research to include T1-based quantitative semi-automated and automated methods for mild cognitive impairment and Alzheimer’s Disease diagnosis, tracking and prognosis (Dale, et al., 1999; Chupin, et al., 2009; Davatzikos, et al., 2009; Leung, et al., 2010; Leung, et al., 2013; Misra, et al., 2016; Schuff, et al., 2009; Sorensen, et al, 2016).

The development of semi-automatic and automatic derivation of resting-state functional connectivity biomarkers for Alzheimer’s Disease classification is comparatively new (Grecius, et al., 2004; Sepulcre, et al., 2017; and Wiepert, et al., 2017). First, Grecius, et al., (2004) applied a goodness-of-fit test between an ICA-derived PCC-based or whole DMN-based templates and individual subject images to classify Alzheimer’s Disease or healthy ageing individuals. Second, Wiepert, et al., (2017), proposed a network failure quotient (NFQ). The measure combined the increases and decreases in DMN subsystem connectivity with Alzheimer’s Disease progression into a single statistic. The authors demonstrated that the statistic (“trained” in an ADNI cohort) had the greatest effect size in differentiating CN and AD in a distinct cohort (Mayo Clinic).

Last, Sepulcre, et al., (2017) applied graph theory methods to demonstrate distinct associations between functional connectivity changes in cognitively normal aging brains and cortical patterns of *in vivo* tau and amyloid β positron emission tomography imaging. A remaining challenge is to fully characterize classification sensitivity and specificity of functional connectivity biomarkers compared to CSF-derived, plasma-derived and *in vivo* and structural imagine modalities.

### 4.5. Conclusion

The contribution of the present study in the context of established structural and emerging resting-state functional connectivity biomarkers is four-fold. First, we demonstrated that functional connectivity changes were detected where structural changes were not yet in evidence using widely available imaging clinic resources. Second, we measured functional connectivity changes anchored in the MTL structures directly implicated in the earliest stages of Alzheimer’s Disease. Third, we observed a complex pattern of changes: monotonic and non-monotonic, and persistent and reverting patterns of hyperconnectivity and hypoconnectivity, in a spatial organization closely related to the default-mode network. Last, we adopted a multi-modal functional-structural approach and demonstrated differential potential clinical value of resting-state functional connectivity biomarkers in Alzheimer’s Disease diagnosis, tracking and prognosis.

## 5. Acknowledgements

Data collection and sharing for this project was funded by the Alzheimer’s Disease Neuroimaging Initiative (ADNI) (National Institutes of Health Grant U01 AG024904) and DOD ADNI (Department of Defense award number W81XWH-12-2-0012). ADNI is funded by the National Institute on Aging, the National Institute of Biomedical Imaging and Bioengineering, and through generous contributions from the following: AbbVie, Alzheimer’s Association; Alzheimer’s Drug Discovery Foundation; Araclon Biotech; BioClinica, Inc.; Biogen; Bristol-Myers Squibb Company; CereSpir, Inc.; Cogstate; Eisai Inc.; Elan Pharmaceuticals, Inc.; Eli Lilly and Company; EuroImmun; F. Hoffmann-La Roche Ltd and its affiliated company Genentech, Inc.; Fujirebio; GE Healthcare; IXICO Ltd.; Janssen Alzheimer Immunotherapy Research & Development, LLC.; Johnson & Johnson Pharmaceutical Research & Development LLC; Lumosity; Lundbeck; Merck & Co., Inc.; Meso Scale Diagnostics, LLC.; NeuroRx Research; Neurotrack Technologies; Novartis Pharmaceuticals Corporation; Pfizer Inc.; Piramal Imaging; Servier; Takeda Pharmaceutical Company; and Transition Therapeutics. The Canadian Institutes of Health Research is providing funds to support ADNI clinical sites in Canada. Private sector contributions are facilitated by the Foundation for the National Institutes of Health (www.fnih.org). The grantee organization is the Northern California Institute for Research and Education, and the study is coordinated by the Alzheimer’s Therapeutic Research Institute at the University of Southern California. ADNI data are disseminated by the Laboratory for Neuro Imaging at the University of Southern California.

## 8. Supplemental Material - Tables

**1. Supplemental Information Table 1 (Cortical Thickness ANOVA Results).** Excel table containing the ANOVA F statistic and *p* value, *post hoc* pair-wise *t* statistic and *p* value, pair-wise Cohen’s *d*, group average, and group average standard error, for 68 cortical region thickness estimates.

**2. Supplemental Information Table 2 (Subcortical Segmentation ANOVA Results).** Excel table containing the ANOVA F statistic and *p* value, *post hoc* pair-wise *t* statistic and *p* value, pair-wise Cohen’s *d*, group average, and group average standard error, for 16 hippocampal subfield volume estimates.

**3. Supplemental Information Table 3 (Hippocampal Subfields ANOVA Results).** Excel table containing the ANOVA F statistic and *p* value, *post hoc* pair-wise *t* statistic and *p* value, pair-wise Cohen’s *d*, group average, and group average standard error, for 22 subcortical structure volume estimates.

**4. Supplemental Information Table 4 (Left Hemisphere ANOVA Raw Results).** Excel tables containing left significant ROI pairs of interest, their ANOVA sample F statistic and *p* value, repeated measures permuted F statistic and *p* value, permutation estimated *post hoc* pair-wise *t* test *p* value, pair-wise Cohen’s *d*, and group averages.

**5. Supplemental Information Table 5 (Right Hemisphere ANOVA Raw Results).** Excel tables containing right significant MCI-CN contrasts, their ANOVA sample F statistic and *p* value, repeated measures permuted F statistic and *p* value, permutation estimated *post hoc* pair-wise *t* test *p* value, pair-wise Cohen’s *d*, and group averages.

**6. Supplemental Information Table 5 (Left and Right Tabulated ANOVA Results).** Excel tables containing right significant MCI-CN and AD-CN and AD-MCI contrasts, their ANOVA sample F statistic and *p* value, repeated measures permuted F statistic and *p* value, permutation estimated *post hoc* pair-wise *t* test *p* value, pair-wise Cohen’s *d*, and group averages, and including MNI coordinates. Tables correspond to Tables 3 and 4 in the main text.

### 9. Supplemental Material - Figures

**Supplemental Figure 1.**
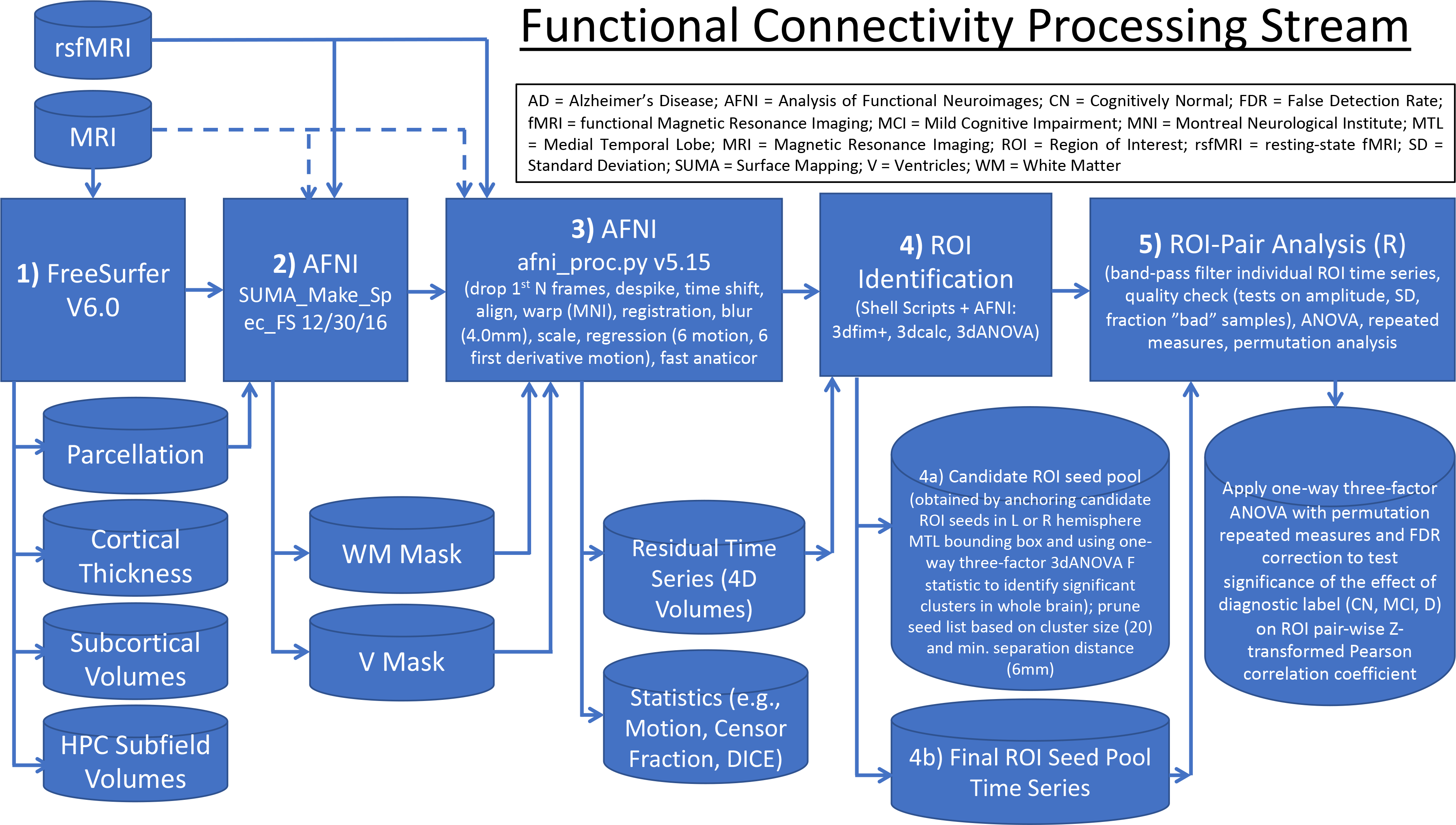
Resting-state fMRI processing block diagram.

**Supplemental Figure 2.**
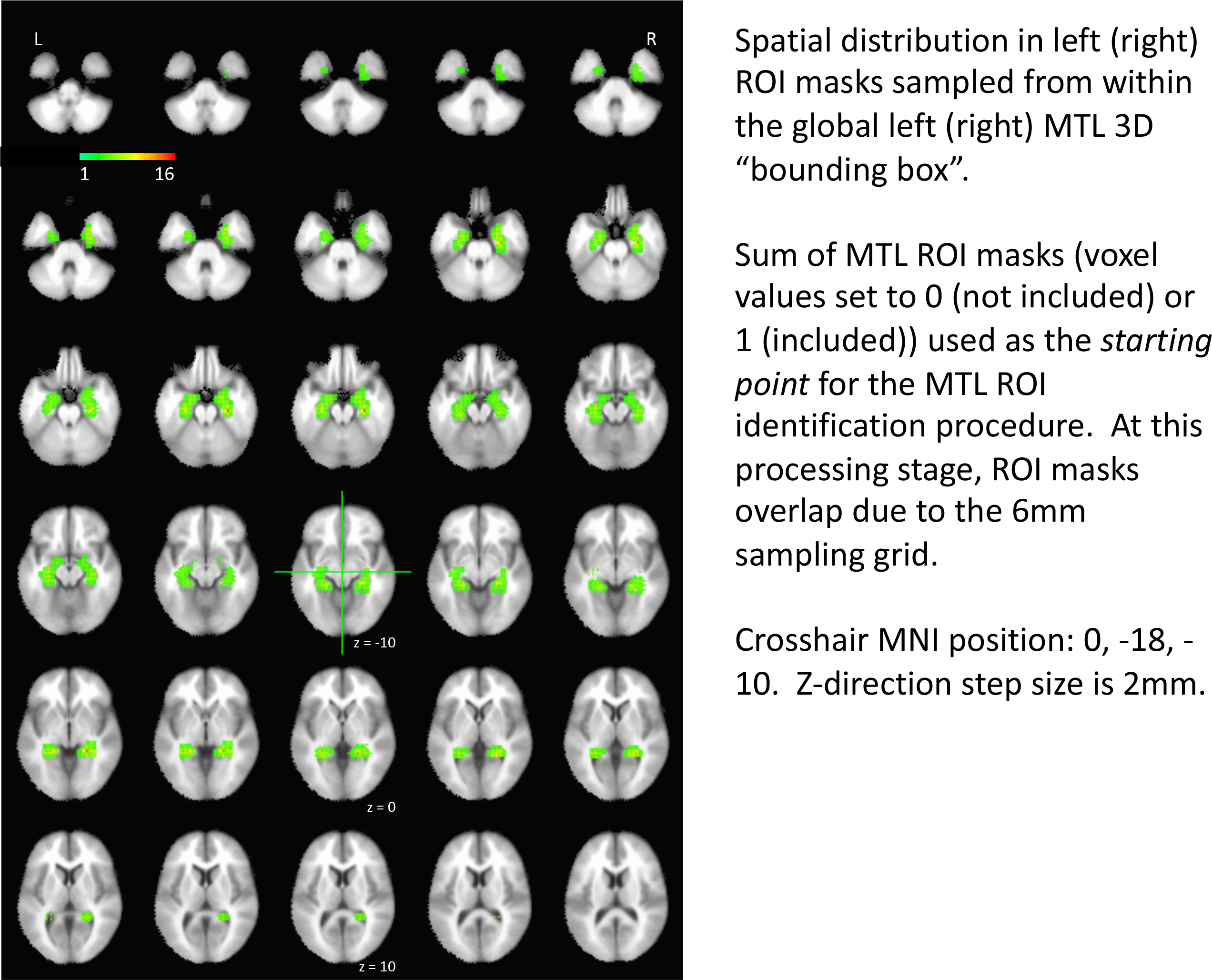
Spatial distribution in left (right) ROI masks sampled from within the global left (right) MTL 3D “bounding box”.

**Supplemental Figure 3.**
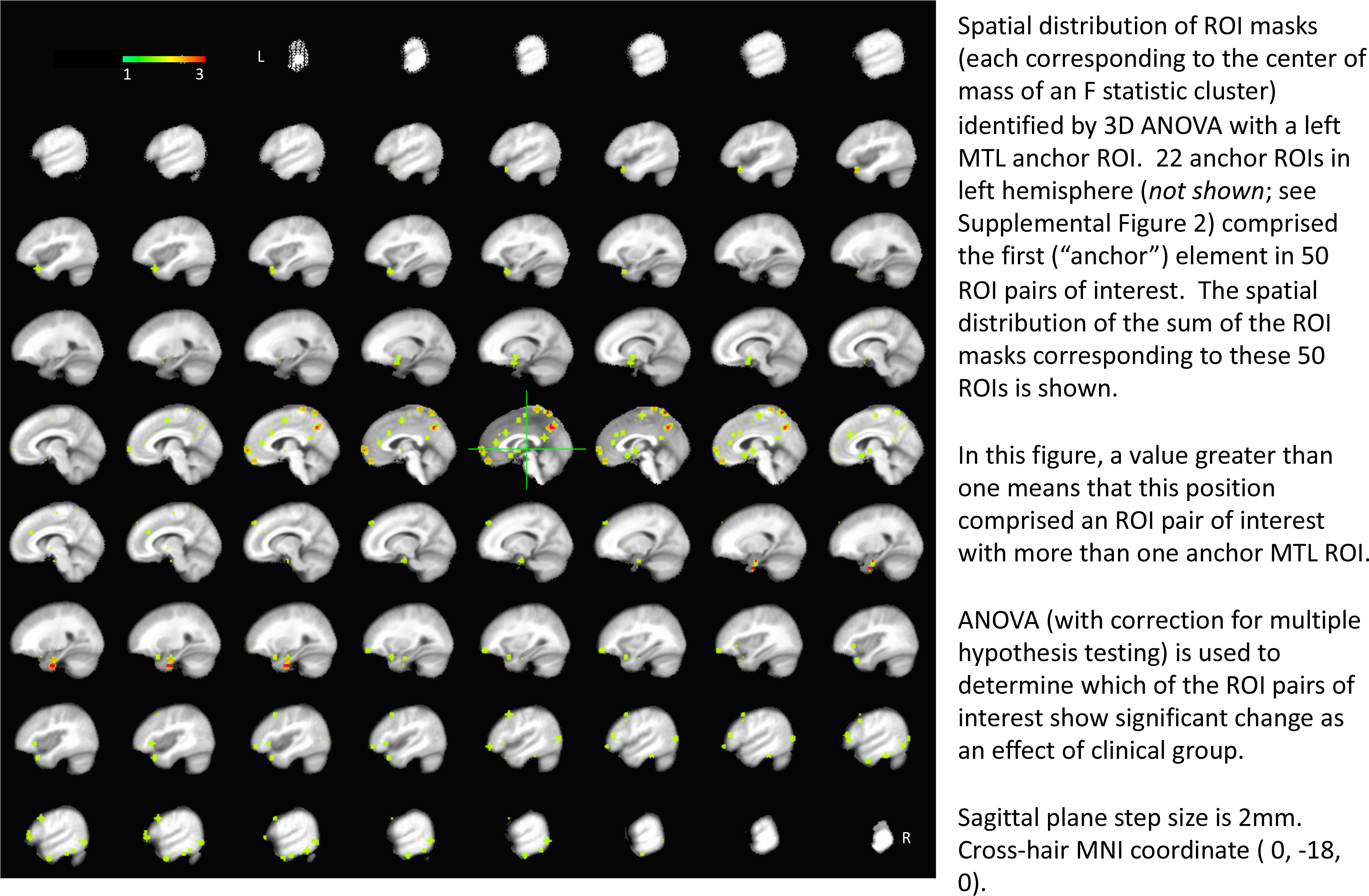
Spatial distribution of ROI masks (each corresponding to the center of mass of an F statistic cluster) identified by 3D ANOVA with a left MTL anchor ROI.

**Supplemental Figure 4.**
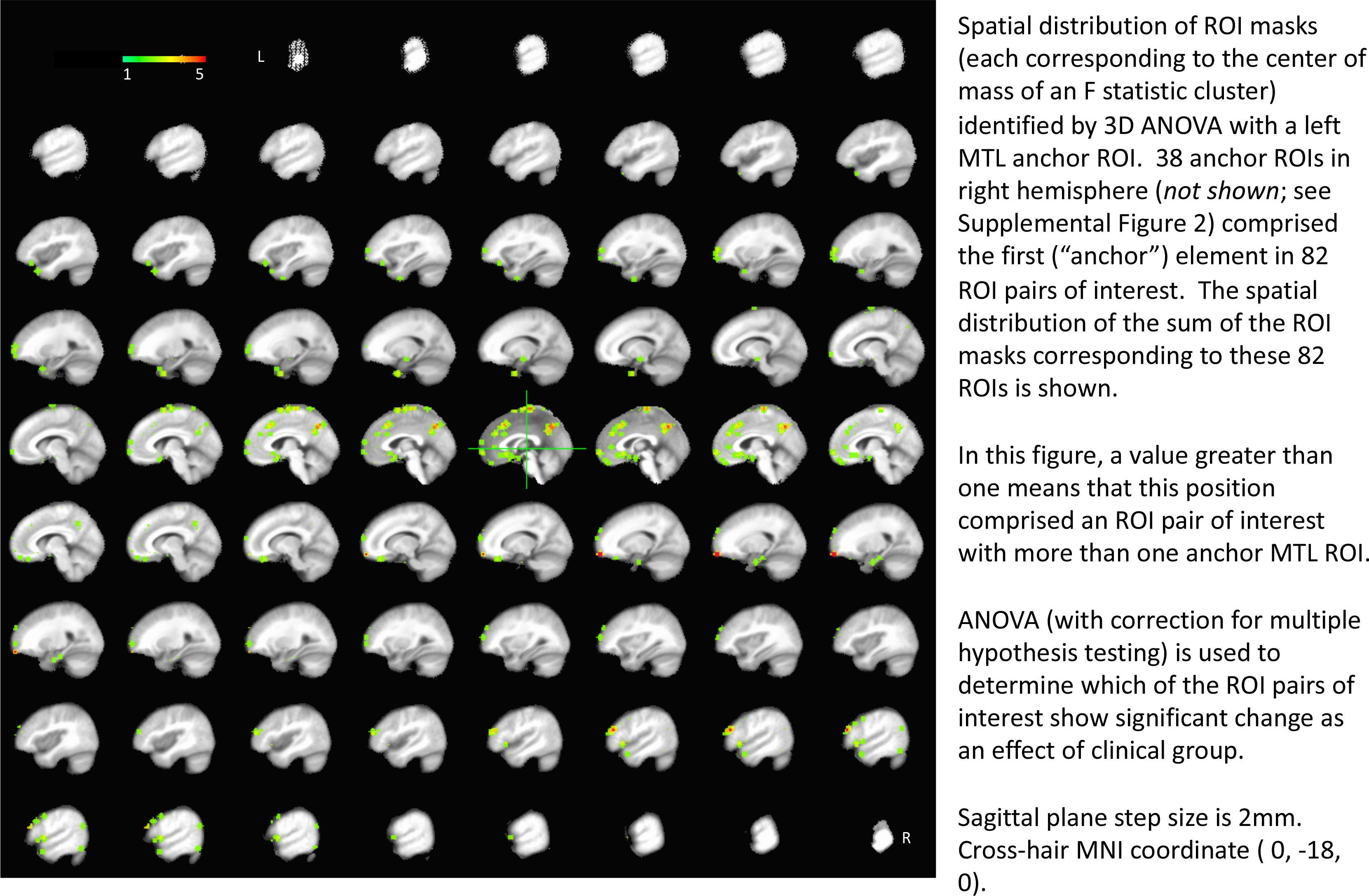
Spatial distribution of ROI masks (each corresponding to the center of mass of an F statistic cluster) identified by 3D ANOVA with a right MTL anchor ROI.

**Supplemental Figure 5.**
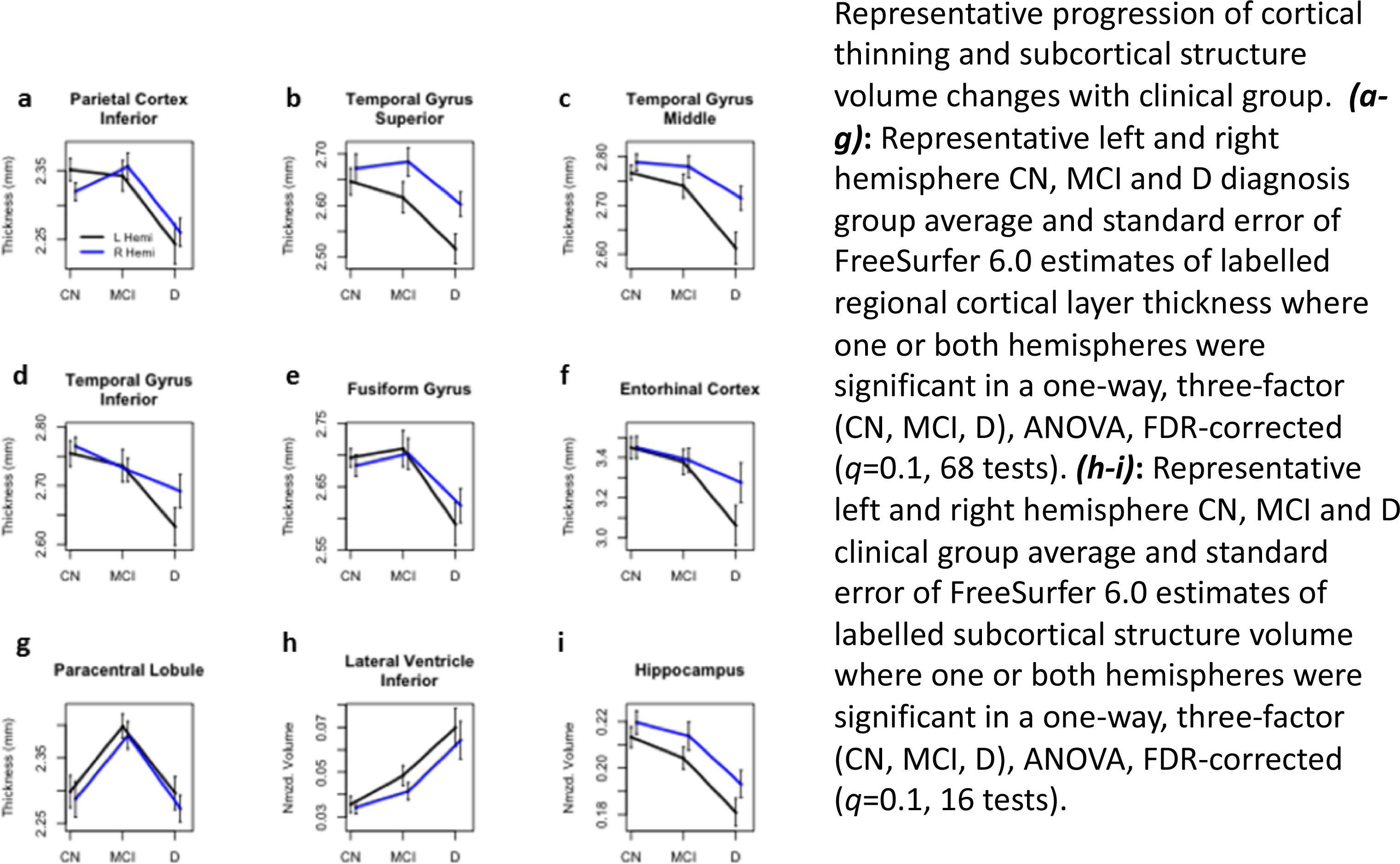
Progression of cortical thinning in representative regions and subcortical structure volume with clinical group. ANOVA results are shown in table format in Supplemental Tables 1-3.

**Supplemental Figure 6.**
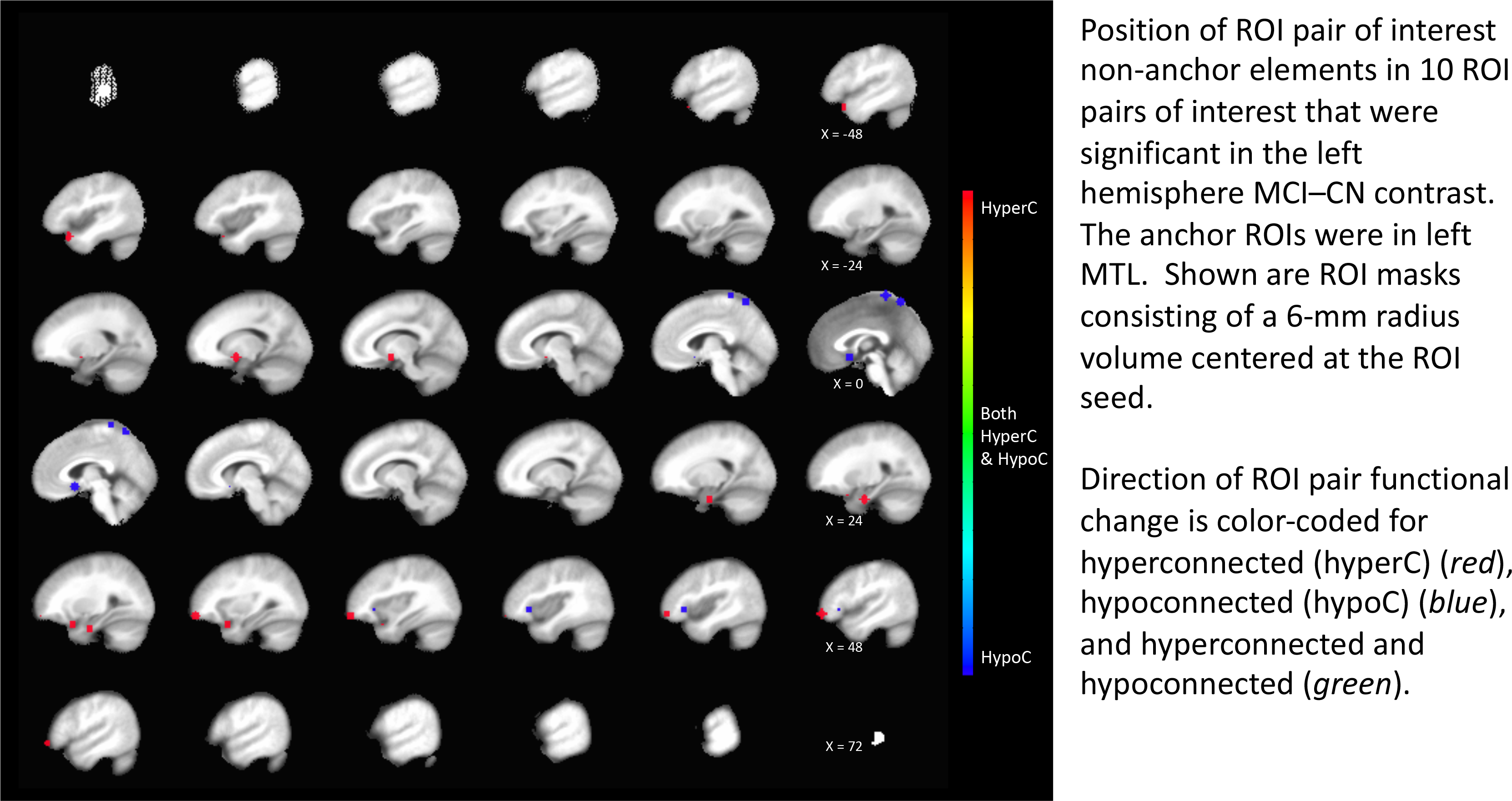
MCI-CN (left MTL anchors). Spatial distribution of the non-anchor ROI masks (where the anchor ROI mask was in left MTL) for the 10 significant ROI pairwise comparisons in the MCI-CN contrast.

**Supplemental Figure 7.**
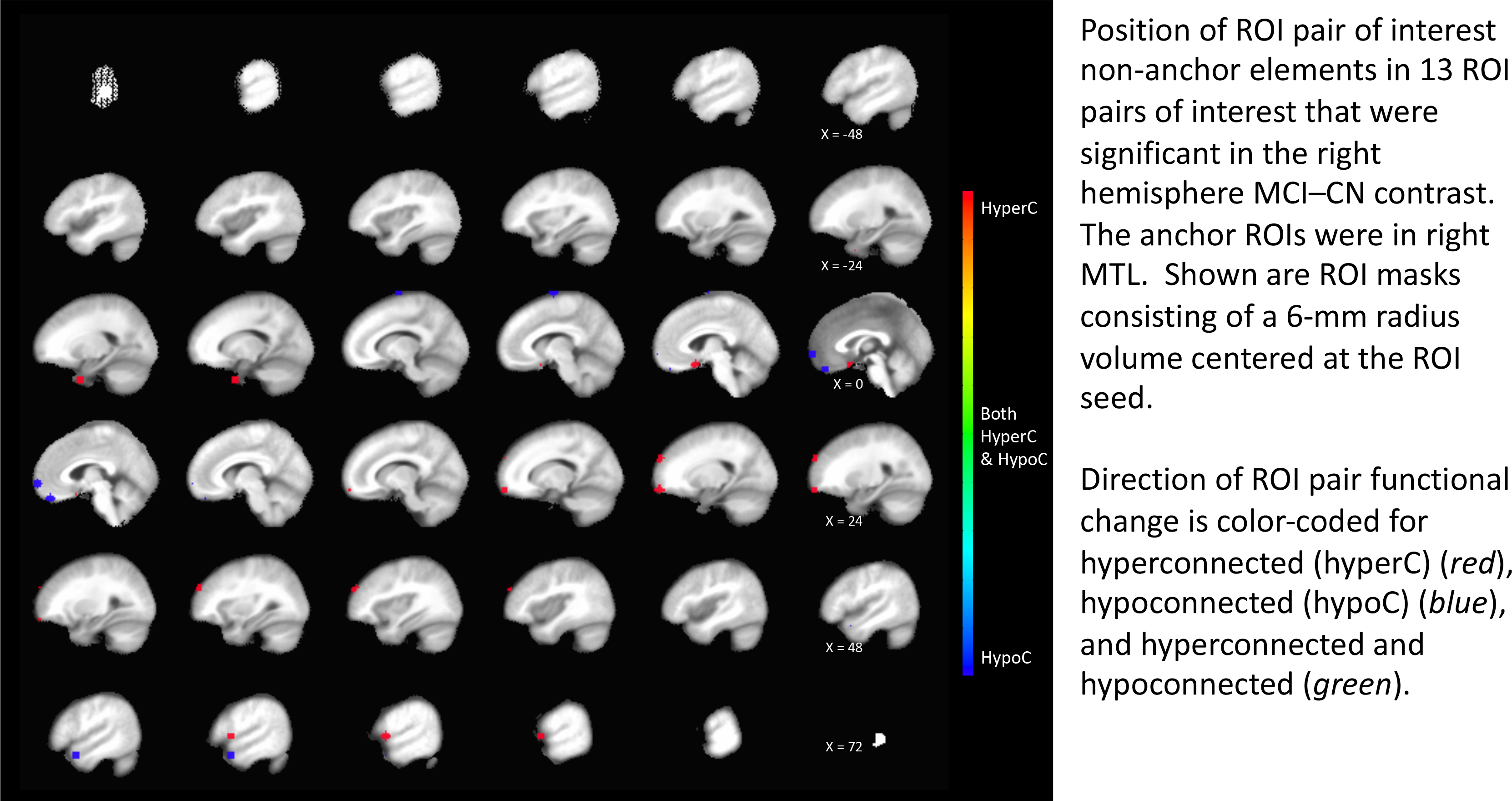
MCI-CN (right MTL anchors). Spatial distribution of the non-anchor ROI masks (where the anchor ROI mask was in right MTL) for the 10 significant ROI pairwise comparisons in the MCI-CN contrast.

**Supplemental Figure 8.**
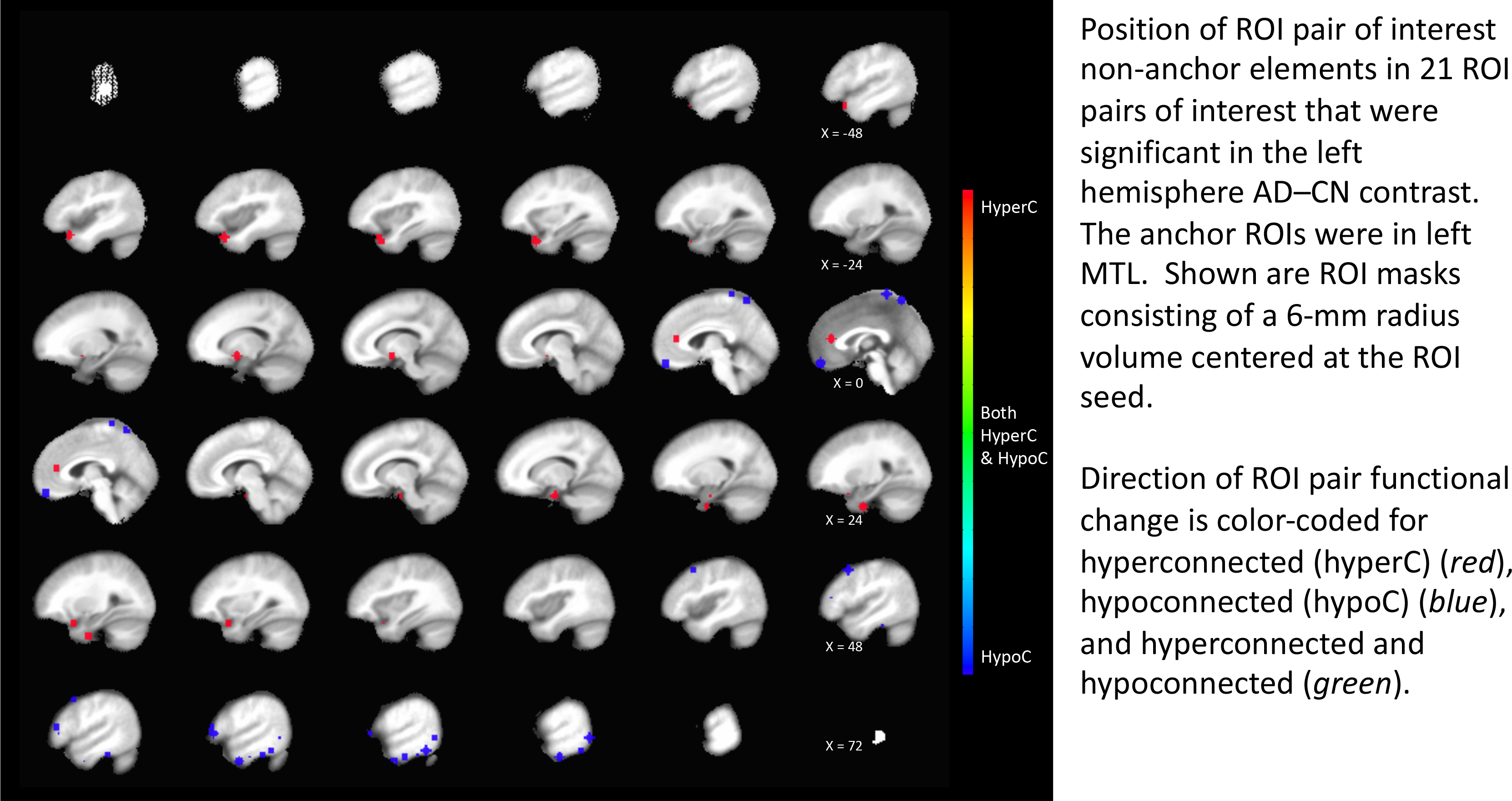
AD-CN (left MTL anchors). Spatial distribution of the non-anchor ROI masks (where the anchor ROI mask was in left MTL) for the 21 significant ROI pairwise comparisons in the AD-CN contrast.

**Supplemental Figure 9.**
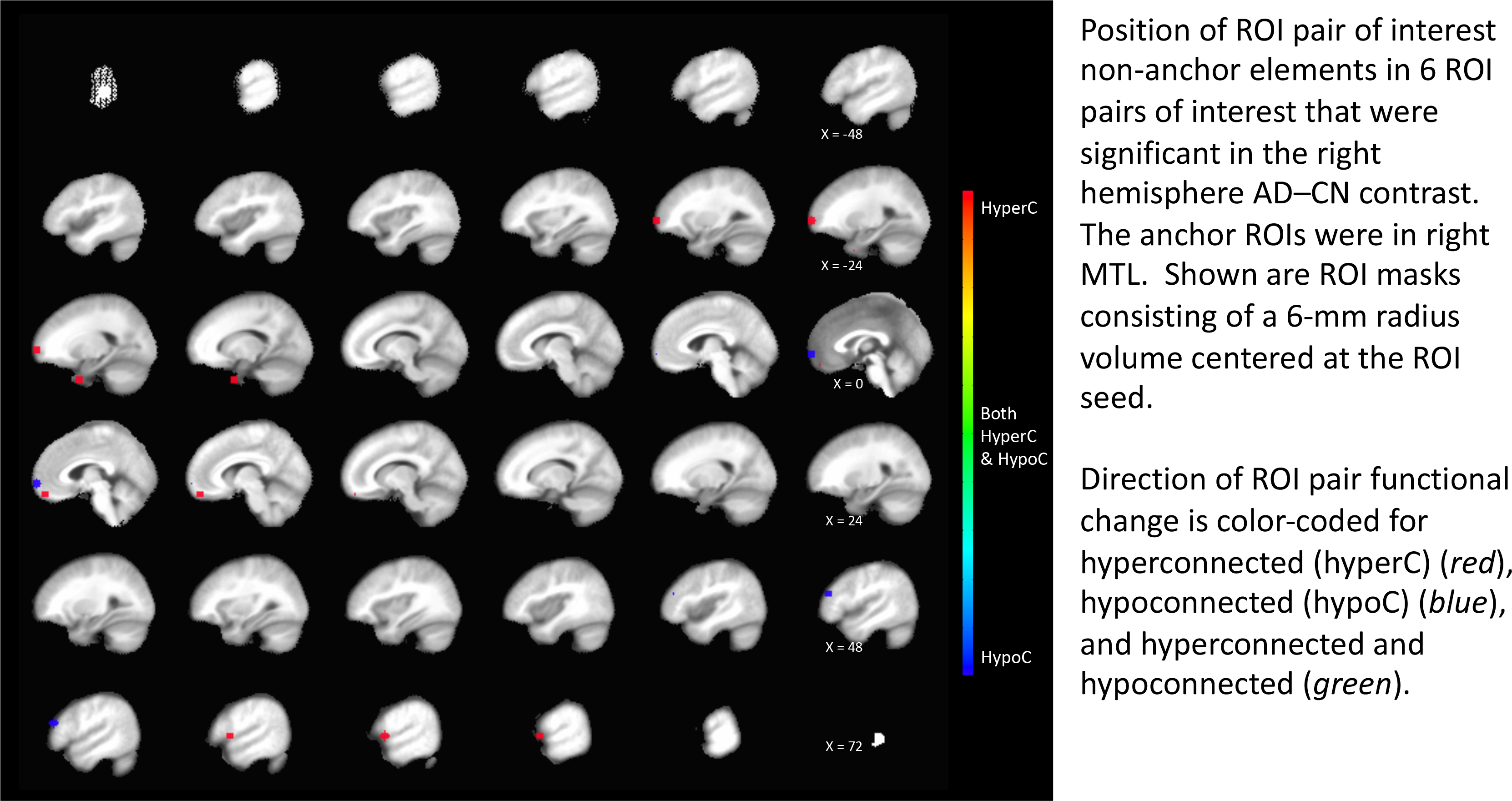
AD-CN (right MTL anchors). Spatial distribution of the non-anchor ROI masks (where the anchor ROI mask was in right MTL) for the 6 significant ROI pairwise comparisons in the AD-CN contrast.

## Footnotes

4 Supplemental Information Table 6 (Worksheet MCI-CN_Detail) provides additional details, including ROI MNI coordinates.

5 Supplemental Information Table 6 (Worksheet MCI-CN_Detail) provides additional details, including ROI MNI coordinates.

## Abbreviations

ADNI =: Alzheimer’s Disease Neuroimaging Initiative
CA =: cornu ammonis
CDR =: clinical dementia rating
CDRSB =: clinical dementia rating, sum of boxes
CDT =: cluster determination threshold
CN=: cognitively normal
CSF =: cerebrospinal fluid
D =: dementia
DMN =: default-mode network
EPI =: echo-planar image
FWHM =: full width half maximum
ICA =: independent components analysis
ICV =: intra-cranial volume
MCI =: mild cognitive impairment
MMSE =: mini-mental state examination
MTL =: medial temporal lobe
PCC =: posterior cingulate cortex
PHG =: parahippocampal gyrus
rsfMRI =: resting-state functional MRI
ROI =: region of interest

